# Small Molecule Agonists of TREM2 Reprogram Microglia and Protect Synapses in Human Alzheimer’s Models

**DOI:** 10.64898/2026.01.19.700278

**Authors:** Hossam Nada, Shaoren Yuan, Farida El Gaamouch, Sungwoo Cho, Moustafa T. Gabr

## Abstract

Triggering receptor expressed on myeloid cells-2 (TREM2) is a key immune receptor in the central nervous system that regulates microglial phagocytosis, survival, and neuroinflammatory responses. TRME2 variants have been established as genetic risk factors for Alzheimer’s disease (AD). However, the therapeutic development of TREM2 modulators has been limited to antibody-based approaches that face limitations in blood-brain barrier penetration and manufacturing scalability. Furthermore, there are no FDA approved TREM2 therapeutics available to date marking an unmet therapeutic gap. Herein, we report the identification of the first TREM2 small molecule submicromolar binders as a result of optimizing compound **4a** to yield **S9** with TREM2 binding affinity of 0.95 µM. **S9** demonstrated robust TREM2 agonism in cellular assays where it induced proximal Syk phosphorylation, activated downstream NFAT transcriptional signaling, enhanced APOE internalization and microglial phagocytic capacity. Pharmacokinetic profiling of the optimized hits revealed **S9** to exhibit improved drug-likeness compared to **4a** with 7-fold enhanced aqueous solubility, superior metabolic stability, reduced intrinsic clearance and a 9-fold improved hERG safety margin. Functional validation in human iPSC-derived microglia confirmed that **S9** suppresses amyloid-beta (Aβ)-induced IL-1β secretion through a TREM2-dependent mechanism. In human neuron-microglia co-culture models exposed to amyloid stress, **S9** treatment preserved synaptic integrity as measured by PSD95 expression that indicates promising neuroprotective activity. Together, these findings establish **S9** as a first-TREM2 submicromolar small molecule TREM2 agonist which is orally bioavailable with favorable pharmacokinetic properties and promising therapeutic potential for the treatment of Alzheimer’s disease.

## 1. Introduction

Triggering receptor expressed on myeloid cells-2 (TREM2) is a transmembrane innate immune receptor that is predominantly expressed on myeloid lineage cells which include dendritic cells, tissue- resident macrophages and microglia within the central nervous system (CNS)^1,2^. TREM2 structure is comprised of an extracellular V-type immunoglobulin (Ig) domain which is linked to a transmembrane helix via a short stalk region and a cytoplasmic tail lacking intrinsic signaling motifs^3,4^. Signal transduction occurs through the transmembrane helix which interacts with the adaptor protein DAP12^5^ (encoded by TYROBP) which contains an immunoreceptor tyrosine-based activation motif (ITAM^6^). Ligand engagement of TREM2 through DAP12 results in a signaling cascade that is responsible for regulating key microglial functions including phagocytosis, survival, proliferation, motility and lysosomal activity^7,8^. Additionally, proteolytic cleavage or alternative splicing generates soluble TREM2 (sTREM2^9^) which retains immunomodulatory functions.

Under physiological conditions, TREM2 plays an essential role in maintaining CNS homeostasis by enhancing microglial-mediated phagocytosis and clearance of apoptotic cells, myelin debris, and misfolded protein aggregates such as amyloid-beta (Aβ), aggregated tau and α-synuclein species^10,11^. TREM2 also modulates microglial polarization, balancing proinflammatory (M1-like) and anti- inflammatory (M2-like) phenotypes to regulate neuroinflammation and support immune surveillance, synaptic pruning, and debris clearance^12,13^. Alzheimer’s disease (AD) represents a major unmet medical challenge, characterized by extracellular Aβ deposits, intracellular neurofibrillary tangles of hyperphosphorylated tau, and sustained neuroinflammation.

TREM2 dysfunction has been established as a key contributor to AD pathogenesis due to the discovery of the associated genetic risk factors of TREM2 variants, particularly R47H, R62H, and T66M. These loss-of-function mutations impair microglial recruitment and clustering around Aβ plaques, resulting in less compact, more diffuse deposits^14,15^. In addition, TREM2 deficiency has been shown to compromise microglial survival^16^ during reactive microgliosis which heightens susceptibility to Aβ- mediated toxicity and aggravates neuronal injury^17,18^. Moreover, Transcriptomic studies have shown that TREM2-deficient microglia fail to acquire protective gene expression programs where increased neuritic degeneration was observed consistently across models^19,20^. Conversely, TREM2 overexpression enhances phagocytic capacity, reduces amyloid burden and neuritic pathology, while agonistic antibodies targeting TREM2 promote plaque compaction, lower Aβ levels, attenuate neuritic injury, and improve cognitive outcomes in preclinical models^21,22^.

Together, these findings highlight TREM2 as a key therapeutic target for AD which has the potential to reverse or slow AD disease progression unlike current AD therapies that only provides symptomatic relief. To date, most TREM2 therapeutic strategies have focused on antibody-based TREM2 modulation which face the inherent challenge of restricted blood-brain barrier (BBB) penetration, high production costs, immunogenicity risks and prolonged systemic half-lives that may extend immune-related adverse events^23^. Toward avoiding these limitations, it is necessary to develop small molecule which can offer superior BBB permeability, oral bioavailability, lower immunogenicity potential, precise pharmacokinetic optimization for adjustable dosing, and reduced manufacturing costs that could improve patient accessibility.

To date there are no FDA approved small molecule TREM2 therapeutics which marks the development of novel small molecule TREM2 modulators as a critical and unmet therapeutic need. Herein, we describe the optimization efforts carried out to optimize our recently identified DEL-screening micromolar TREM2 hit (compound **4a**^24^) to yield ten novel derivatives with submicromolar direct TREM2 agonists. In addition, the optimized derivatives show better in-vitro pharmacokinetics and ability to improve AD related outcomes when tested in AD model.

## 2. Results and Discussion

### 2.1. Chemistry

The thiophene ring was selected as the focus for optimization and was replaced with benzene- based derivatives, yielding compounds **S1-10**. The room temperature for 6 hours to afford the synthetic route commenced with the bromination of 3-amino-4-methoxy-N-methylbenzamide (1) using N-bromosuccinimide (NBS) and hydrochloric acid (HCl) at brominated intermediate (2). Subsequently, compound **2** was subjected to amide coupling with 3-methylenecyclobutane-1-carboxylic acid in the presence of 1-ethyl-3-(3-dimethylaminopropyl)carbodiimide hydrochloride (EDC·HCl) and 4- dimethylaminopyridine (DMAP) at room temperature for 2 hours, which furnished intermediate 3. Finally, a Suzuki-Miyaura cross-coupling reaction was employed to introduce diverse aryl substituents at the brominated position, generating the final compounds **S1-10**.

**Scheme 1.**
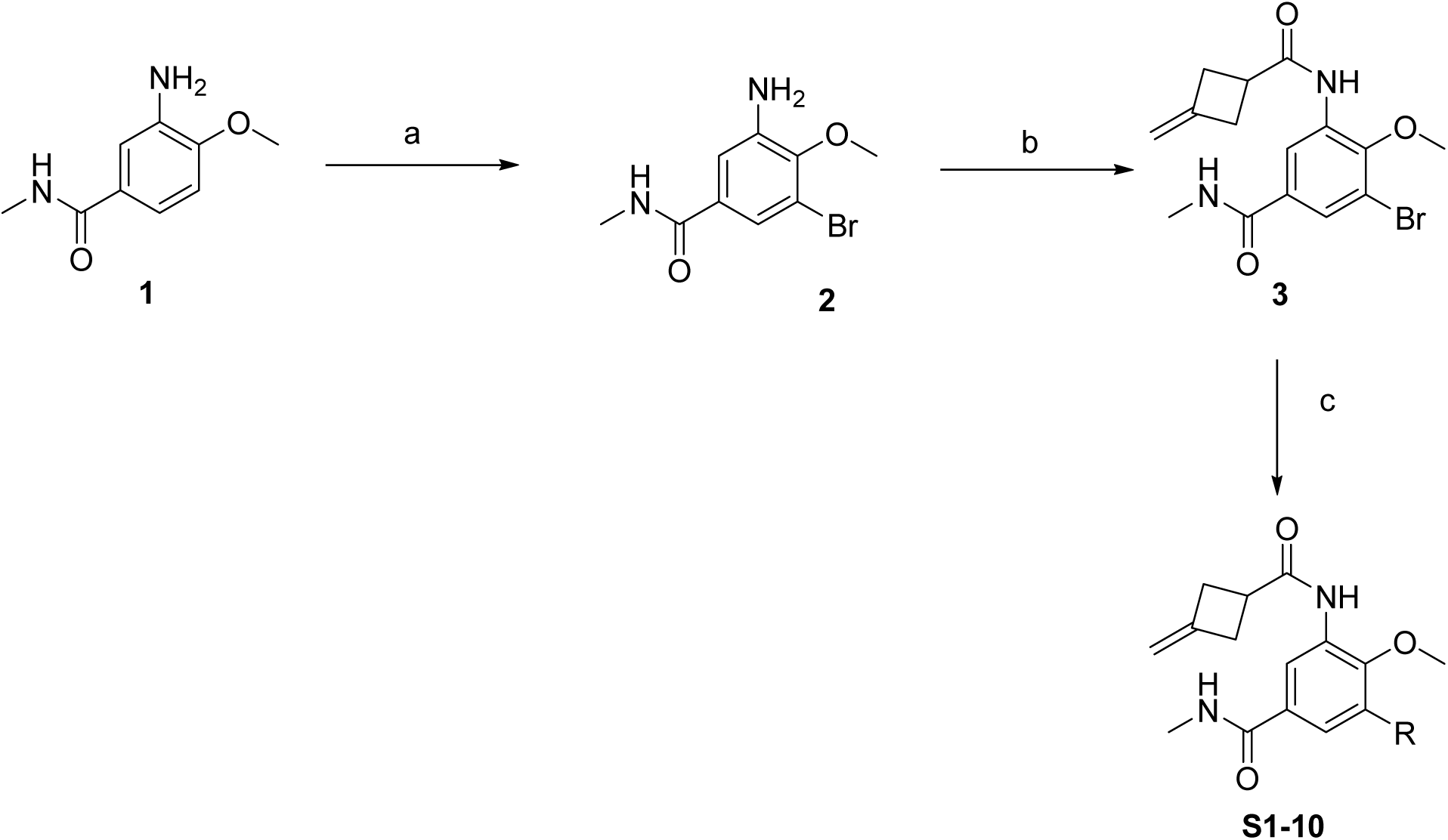
Synthesis of compounds S1-10. (**a**) 3-Sulfolene, NBS, HCL, rt, 6h; (**b**) 3-methylenecyclobutane-1-carboxylic acid, EDC·HCl, DMAP, rt, 2h; (**c**) Boronic acid derivatives DME/water (5:1), Pd(PPh_3_)_4_, Cs_2_CO_3_, 85 °C, Overnight.

### 2.2. TRIC-based affinity screening

To evaluate the binding affinity of synthesized derivatives of compound 4a, a previously reported TREM2 agonist, to TREM2, we performed biophysical screening using Temperature-Related Intensity Change (TRIC) technology (**Figure 2**). All test compounds were screened at a concentration of 30 μM against purified TREM2 protein. The normalized fluorescence intensity (Fnorm, ‰) was measured for each compound, and hits were defined as compounds exhibiting Fnorm values exceeding five standard deviations from the negative control mean. Among the investigated compounds, six derivatives (**S2**, **S5**, **S7**, **S8**, **S9**, and **S10**) were identified as TREM2 binders, displaying Fnorm values comparable to the positive control (compound **4a**). The remaining derivatives showed Fnorm values within the range of the negative control, indicating no detectable binding to TREM2 under these assay conditions. These results suggest that specific structural modifications of the **4a** scaffold can maintain or enhance TREM2 binding activity.

**Figure 1.**
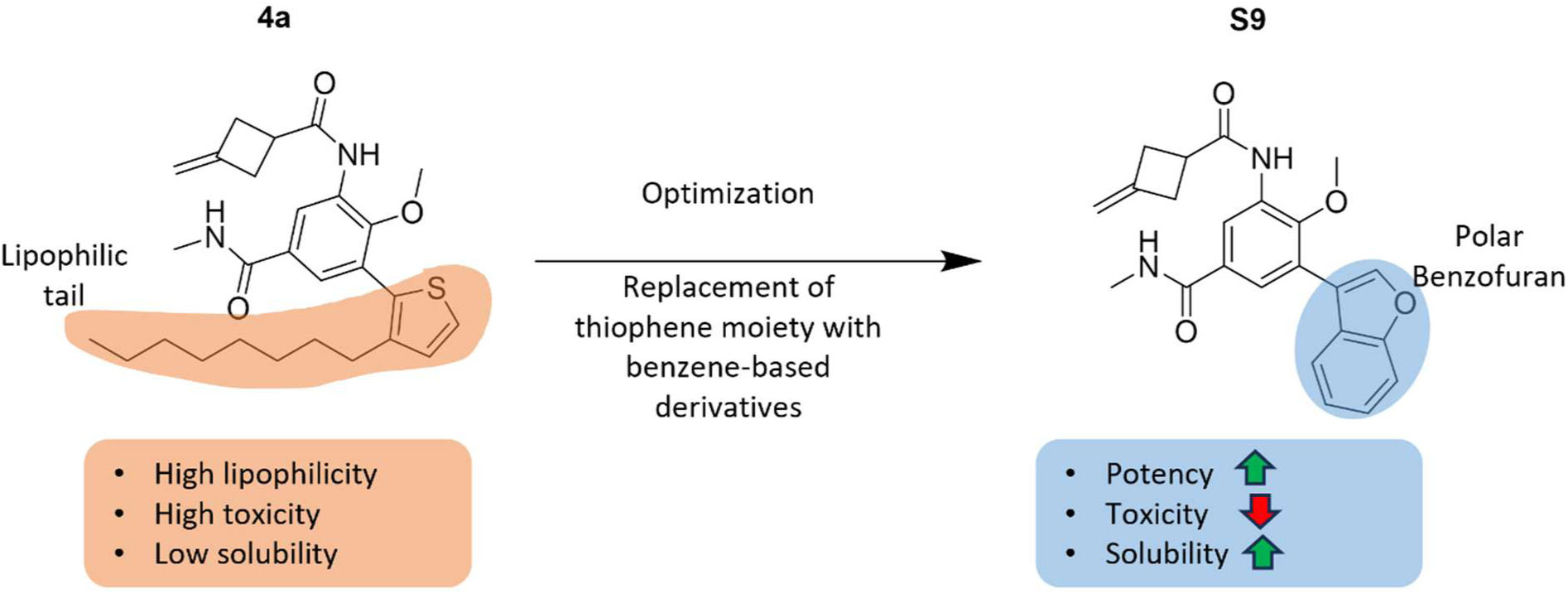
Structure-based optimization of TREM2 agonist 4a to lead compound S9. The key structural modification involved replacement of the lipophilic thiophene-alkyl chain (highlighted in orange) with a polar benzofuran moiety (highlighted in blue), resulting in substantial improvements in drug-like properties and potency.

**Figure 2.**
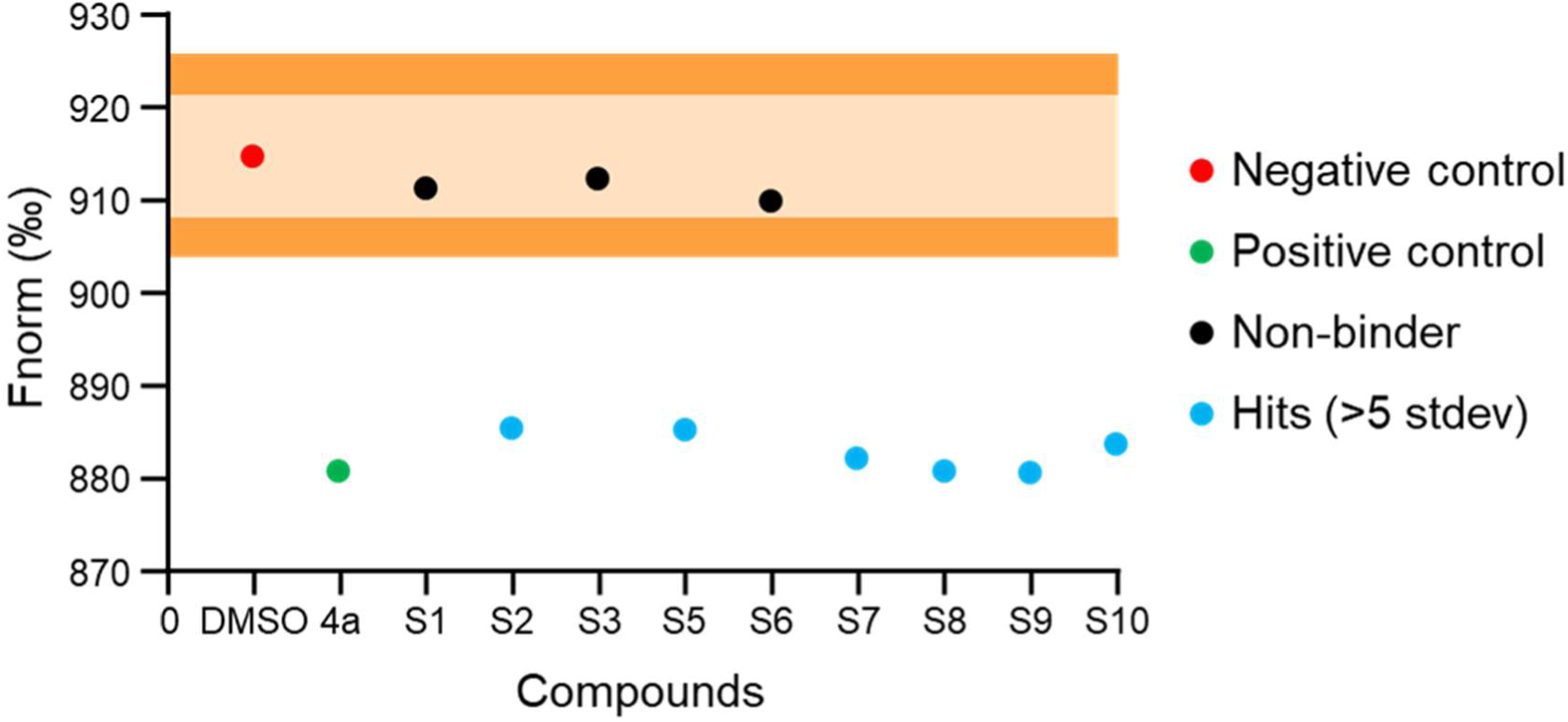
TRIC-based biophysical screening identifies benzene-based derivatives of TREM2 binding compound 4a. Normalized fluorescence intensity (Fnorm, ‰) values from Temperature-Related Intensity Change (TRIC) analysis of synthesized 4a derivatives (30 μM) against TREM2. The dark orange shaded region represents the mean ± 5 standard deviations of the negative control. Red circle, negative control (DMSO vehicle); green circle, positive control (parent compound 4a); black circles, non-binders; cyan circles, hit compounds (>5 SD from negative control). 6 compounds (**S2**, **S5**, **S7**, **S8**, **S9**, and **S10**) were identified as TREM2 binders based on significant deviation from the negative control threshold.

### 2.3. Dose-Dependent TREM2 Binding

Hits that exhibited a binding affinity in the single-dose screening were subjected to a dose-dependent assay (**Figure 3**) against TREM2 using microscale thermophoresis assay (MST). Among the six hit compounds, S5 exhibited no concentration-dependent change in Fnorm values, indicating that this compound was a false positive from the primary screening (**Figure 3B**). In contrast, the remaining five compounds (S2, S7, S8, S9, and S10) demonstrated clear dose-dependent binding curves with TREM2, confirming their identity as genuine TREM2 binders (**Figure 3A, C–F**). The binding affinities (K_D_) were determined as follows: **S2**, 50.91 ± 19.22 μM; **S7**, 1.29 ± 0.15 μM; **S8**, 0.97 ± 0.19 μM; **S9**, 0.95 ± 0.13 μM; and **S10**, 1.42 ± 0.27 μM. Notably, S8 and S9 exhibited the highest binding affinities with sub-micromolar K_D_ values. These results demonstrate that structural optimization of the **4a** scaffold successfully yielded potent TREM2 binders with enhanced binding affinities.

**Figure 3.**
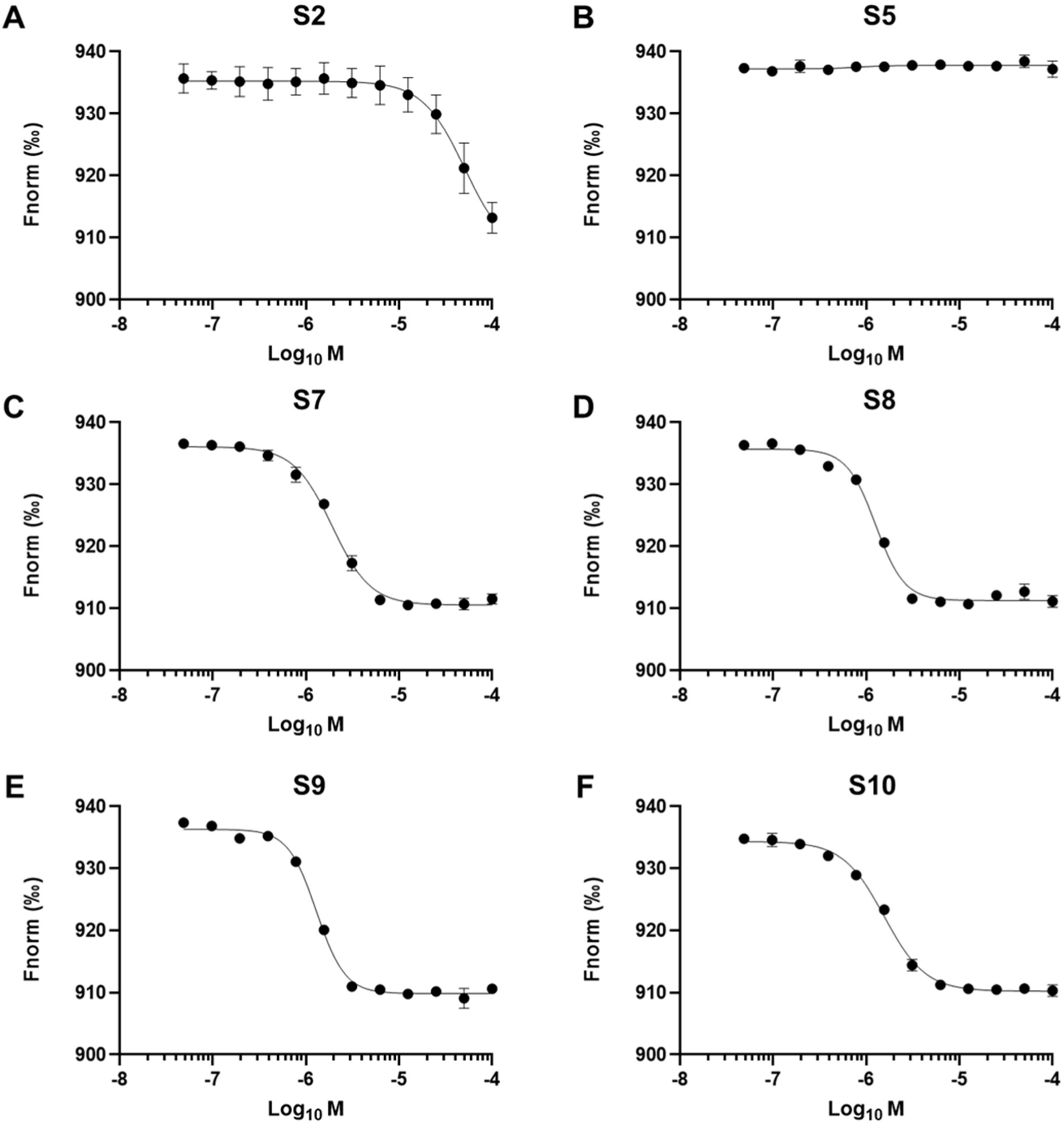
Dose-response analysis of hit compounds binding to TREM2 by Microscale Thermophoresis (MST). MST binding curves of hit compounds identified from TRIC screening against TREM2. Normalized fluorescence intensity (Fnorm, ‰) was plotted as a function of compound concentration (Log_10_ M). (A) S2. (B) **S5**, no detectable binding (false positive). (C) **S7**. (D) **S8**. (E) **S9**. (F) **S10**. Data are presented as mean ± SD from three independent experiments (n=3).

### 2.4. Cell-based assay

Next, the synthesized hit compounds that demonstrated clear dose-response binding profiles were advanced to *in vitro* cellular assays to evaluate their functional activity downstream of receptor engagement. We first assessed their ability to modulate spleen tyrosine kinase (Syk) phosphorylation, a canonical early signaling event following TREM2 activation (**Figure 4A**). Among the investigated hits, compounds **S7** and **S9** induced a robust and statistically significant increase in Syk phosphorylation in a dose-dependent manner. Notably, **S7** produced the strongest effect, closely followed by **S9**, indicating that these compounds function as potent TREM2 agonists capable of efficiently triggering proximal signaling cascades. In contrast, **S8** elicited a more moderate but reproducible dose-dependent enhancement of Syk phosphorylation, suggesting partial agonist activity or reduced signaling efficacy relative to **S7** and **S9**.

**Figure 4.**
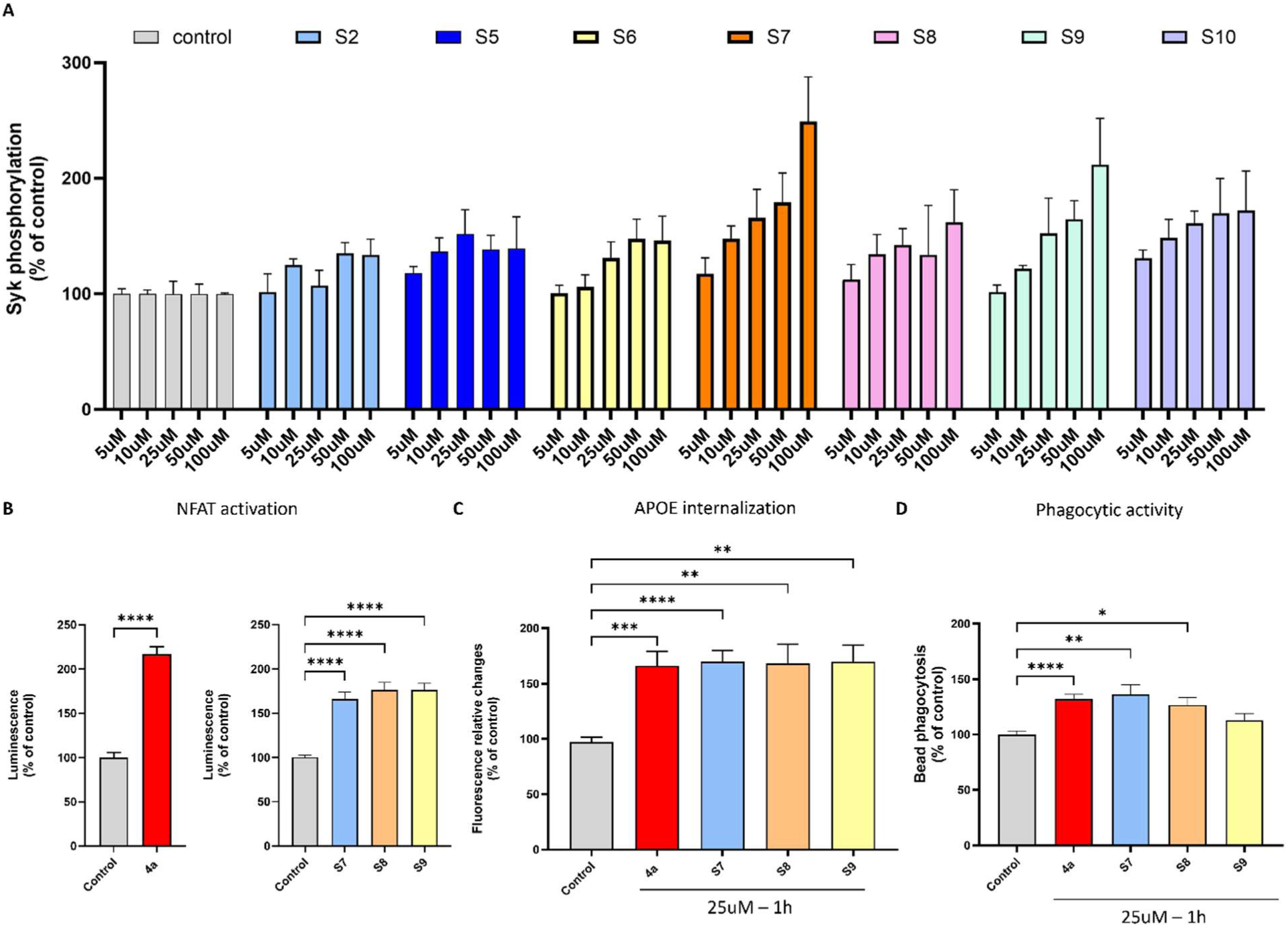
In vitro evaluation of TREM2 engagement and functional activity. (A) Quantification of phospho-SYK levels by AlphaLISA in untreated cells or following treatment with **S2-10** at increasing concentrations (5–100 μM). Phospho-SYK levels are expressed as a percentage of untreated control (N=4). (B) Changes in luciferase signal measured after substrate addition under untreated conditions or following treatment with compounds **4a, S7, S8,** or **S9**. Luminescence is expressed relative to control (N=3). (C) Effect of compounds **4a, S7, S8,** and **S9** (25μM) on APOE internalization after 1h of treatment (N=3). (D) Percentage of BV2 cells containing at least one internalized fluorescent latex bead following treatment with **4a, S7, S8,** or **S9**, expressed relative to control (N = 3; n = 24 images per group). Statistical analyses were performed using Brown-Forsythe and Welch ANOVA with Dunn’s post hoc multiple comparisons. Data are presented as mean ± SEM. *p < 0.05, **p < 0.005, ****p < 0.0001.

Based on these findings, compounds **S7, S8**, and **S9** were selected for additional functional analyses, including evaluation of the specificity of TREM2-dependent SYK phosphorylation. As shown in supplementary material (**Figure S1**), treatment with compounds **S7, S8**, and **S9** resulted in increased SYK phosphorylation in HEK cells expressing TREM2/DAP12, whereas no such effect was observed in parental HEK cells lacking TREM2/DAP12 expression. When compared the three hits against the lead compound **4a** where all three derivatives exhibited comparable levels of NFAT activation (approximately 170%) **(Figure 4B)**, demonstrating that the observed proximal Syk phosphorylation translated into productive downstream transcriptional signaling. This concordance across signaling tiers supports the conclusion that these compounds engage TREM2 in a manner that preserves signal propagation rather than inducing biased or abortive activation.

Importantly, the functional relevance of this signaling was further substantiated by cellular phenotypic assays. Significant enhancements in APOE internalization **(Figure 4C)** and phagocytic activity (**Figure 4D**) were observed across the lead series, indicating that these compounds do not merely bind TREM2 but actively drive therapeutically relevant microglial behaviors. In particular, the ability of **4a, S7,** and **S9** to promote the uptake of amyloid-associated proteins such as APOE, coupled with increased bead phagocytosis, suggests that these agonists may enhance microglial clearance mechanisms implicated in Alzheimer’s disease pathology. Collectively, these findings establish that optimized derivatives of the lead scaffold retain functional efficacy across multiple signaling and phenotypic readouts, reinforcing their potential as small-molecule TREM2 agonists capable of modulating disease-relevant immune functions.

### 2.5. In-vitro Pharmacokinetics

To complement the improved TREM2 binding affinity observed for the second-generation compounds, we performed an evaluation of key in vitro PK parameters for the first-generation lead **4a** and the optimized analogs **S7** and **S9**. This study was intended to enable a side-by-side comparison of developability-relevant properties and to assess whether affinity enhancement was accompanied by improvements in physicochemical and ADME profiles. As summarized in Table 1, compound **4a** exhibited a high lipophilicity (LogD_7.4_ = 6.4), consistent with its long hydrophobic substituent, resulting in poor aqueous solubility (10.6 µM in PBS, pH 7.4) and extensive plasma protein binding (>99.8%). While high lipophilicity translated into favorable passive permeability in the PAMPA assay (2.2 × 10⁻⁵ cm/s), it was also associated with rapid microsomal turnover and high intrinsic clearance, particularly in mouse microsomes (CL_int_ = 68 mL/min/mg), leading to short plasma half-lives. In addition, **4a** showed a higher hERG liability (IC_50_ = 3.2 µM), consistent with its elevated lipophilicity.

**Table 1.** Assessment of the in vitro PK properties of 4a, S7, and S9.^a^.

| Compound | **4a** | **S7** | **S9** |
| --- | --- | --- | --- |
| **LogD<sub>7.4</sub>** | 6.4 | 4.7 | 4.2 |
| **Kinetic solubility ($\mu\text{M}$, PBS pH 7.4)** | 10.6 | 29.5 | 74.1 |
| **PAMPA permeability (cm/s)** | $2.2 \times 10^{-5}$ | $8.5 \times 10^{-6}$ | $3.6 \times 10^{-6}$ |
| **Human PPB (%)** | 99.8 | 99.1 | 98.6 |
| **Mouse plasma $t_{1/2}$ (h)** | 0.6 | 1.8 | 2.4 |
| **Human plasma $t_{1/2}$ (h)** | 0.9 | 2.6 | 3.4 |
| **Mouse microsomal $t_{1/2}$ (h)** | 0.35 | 0.95 | 1.3 |
| **Human microsomal $t_{1/2}$ (h)** | 0.55 | 1.4 | 1.9 |
| **Intrinsic clearance – Mouse (mL/min/mg)** | 68 | 24 | 16 |
| **Intrinsic clearance – Human (mL/min/mg)** | 42 | 14 | 9 |
| **hERG $\text{IC}_{50}$ ($\mu\text{M}$)** | $3.2 \pm 0.4$ | $12.5 \pm 0.9$ | $28.1 \pm 2.7$ |

<sup>a</sup>Values are mean ± SD (n=3).

In contrast, the second-generation compounds **S7** and **S9** displayed a more balanced physicochemical profile (**Table 1**, **Figure 5**). Both compounds showed moderate LogD_7.4_ values (4.7 and 4.2, respectively), which translated into substantially improved kinetic solubility (29.5 µM for **S7** and 74.1 µM for **S9**) while maintaining acceptable passive permeability. Although PAMPA permeability was modestly reduced relative to **4a**, values for both analogs remained within a range compatible with cellular activity.

**Figure 5.**
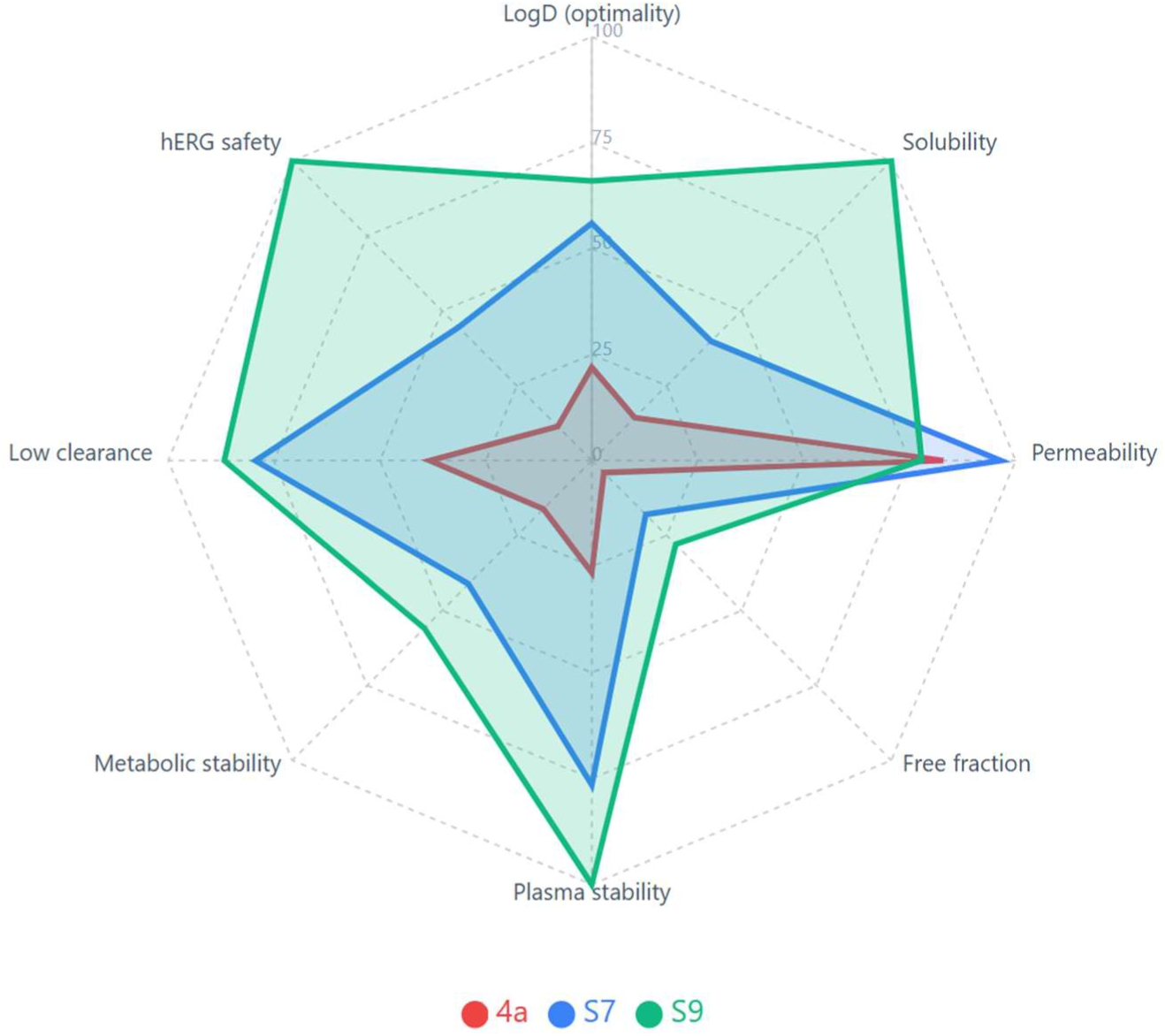
Radar plot comparison of drug-likeness properties for compounds 4a, S7, and S9. Eight key pharmacokinetic and safety parameters were normalized to a 0-100 scale, where higher values indicate more favorable drug-like characteristics.

Importantly, the optimized compounds exhibited marked improvements in metabolic stability. Microsomal half-lives increased approximately three- to five-fold relative to **4a** in both mouse and human systems, accompanied by lower intrinsic clearance values, particularly for **S9** (mouse CL_int_ = 16 mL/min/mg; human CL_int_ = 9 mL/min/mg). These changes translated into longer plasma half-lives, suggesting improved systemic exposure potential. Among the second-generation analogs, **S9** consistently displayed the most favorable PK profile, combining the highest solubility, lowest intrinsic clearance, and lowest hERG liability (IC_50_ = 28 µM). The increased polarity and heteroatom content of **S9** likely contribute to this improved balance, without compromising permeability to an extent that would preclude cellular efficacy.

Taken together, the PK analysis suggests that optimization from **4a** to **S7** and **S9** not only enhanced TREM2 binding affinity but also resulted in a substantially improved developability profile. While **4a** remains a useful first-generation benchmark, its high lipophilicity and metabolic liabilities may limit further progression. In contrast, **S7** and especially **S9** offer a more favorable balance between potency, solubility, metabolic stability, and safety-related liabilities, supporting their selection for further in vivo evaluation. Overall, compound **S9** emerged as the lead compound with the most favorable drug-likeness profile across nearly all assessed parameters

### 2.6. Biological evaluation

To determine whether improvements in TREM2 binding affinity and PK properties translated into enhanced biological activity, we evaluated the effects of the first-generation lead **4a** and the optimized analogs **S7** and **S9** in a human iPSC-derived microglial model of amyloid stress. Microglia were challenged with Aβ_1-42_ oligomers in the presence of increasing concentrations of each compound (1, 5, and 10 µM), and inflammatory output was assessed by quantifying IL-1β secretion. As shown in **Figure 6A**, Aβ oligomer exposure induced a robust increase in IL-1β release relative to vehicle-treated controls. Treatment with **4a** resulted in a modest, dose-dependent reduction in IL-1β levels; however, significant suppression was observed only at the highest concentration tested. In contrast, both **S7** and **S9** produced a more pronounced attenuation of Aβ-induced IL-1β secretion, with measurable effects observed at lower concentrations (1 and 5 µM). Across all tested doses, the second-generation compounds consistently outperformed **4a**, indicating enhanced efficacy in modulating microglial inflammatory responses.

**Figure 6.**
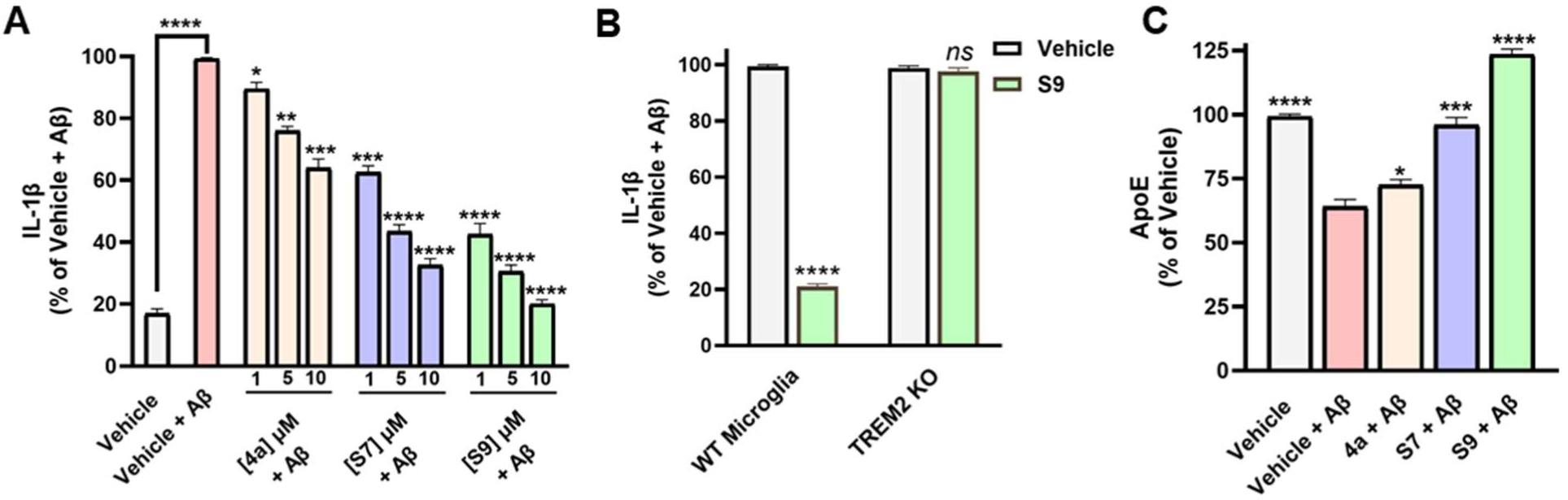
Second-generation TREM2 agonists modulate Aβ-induced microglial responses in a TREM2-dependent manner. **(A)** IL-1β secretion in human iPSC-derived microglia following Aβ_1-42_ oligomer challenge in the presence of **4a**, **S7**, or **S9** (1, 5, and 10 µM). Data are normalized to the Aβ-treated vehicle control and expressed as percent change. **(B)** Loss of **S9**-mediated suppression of IL-1β secretion in TREM2 knockout (KO) microglia confirms TREM2- dependent activity. **(C)** Modulation of secreted ApoE levels in human microglia following Aβ exposure and compound treatment (10 µM). ApoE levels are normalized to vehicle-treated controls. Data are presented as mean ± SD (n = 5). Statistical significance was determined by one-way or two-way ANOVA with Dunnett’s post hoc test. *ns*, not significant; *p* < 0.05 (*), *p* < 0.01 (**), *p* < 0.001 (***), and *p* < 0.0001 (****).

Notably, **S9** exhibited the greatest reduction in IL-1β secretion, suggesting that the affinity enhancement achieved during optimization were accompanied by improved functional activity in a disease-relevant inflammatory context. These findings support a model in which stronger TREM2 engagement leads to more effective suppression of maladaptive microglial activation under amyloid stress.

To confirm that the observed anti-inflammatory effects were TREM2-dependent, we next evaluated compound activity in TREM2 knockout (KO) iPSC-derived microglia. Under Aβ challenge, **S9** significantly reduced IL-1β secretion in wild-type microglia; however, this effect was markedly attenuated in TREM2 KO cells (**Figure 6B**). The loss of compound activity in the absence of TREM2 provides direct genetic evidence that suppression of Aβ-induced inflammatory signaling by **S9** is on-target and TREM2- mediated, rather than resulting from nonspecific anti-inflammatory effects.

Given the central role of the TREM2-APOE axis in disease-associated microglial states, we next examined whether compound treatment altered secreted ApoE levels in Aβ-challenged human microglia. ApoE concentrations in culture supernatants were quantified by ELISA following compound treatment. As shown in **Figure 6C**, treatment with second-generation compounds resulted in a significant modulation of ApoE secretion relative to Aβ-treated vehicle controls. **S7** and **S9** produced a more pronounced effect compared with **4a**, with **S9** again showing the strongest response. These data indicate that optimized TREM2 agonists influence not only inflammatory cytokine output but also a key lipid-associated pathway linked to microglial function in AD.

We next assessed whether the improved microglial modulation observed with **S7** and **S9** translated into functional protection of neuronal synapses. Human iPSC-derived neurons were co-cultured with microglia and exposed to Aβ oligomers in the presence of each compound. Synaptic integrity was quantified by measuring PSD95 puncta density, expressed as a normalized quantitative output. As shown in **Figure 7**, Aβ treatment led to a substantial reduction in PSD95 levels compared with untreated controls, consistent with synaptotoxic effects. Administration of **4a** resulted in partial rescue at higher concentrations, whereas **S7** and **S9** produced significantly greater preservation of PSD95 across multiple concentrations. Among the tested compounds, **S9** consistently demonstrated the most robust synaptic protection.

**Figure 7.**
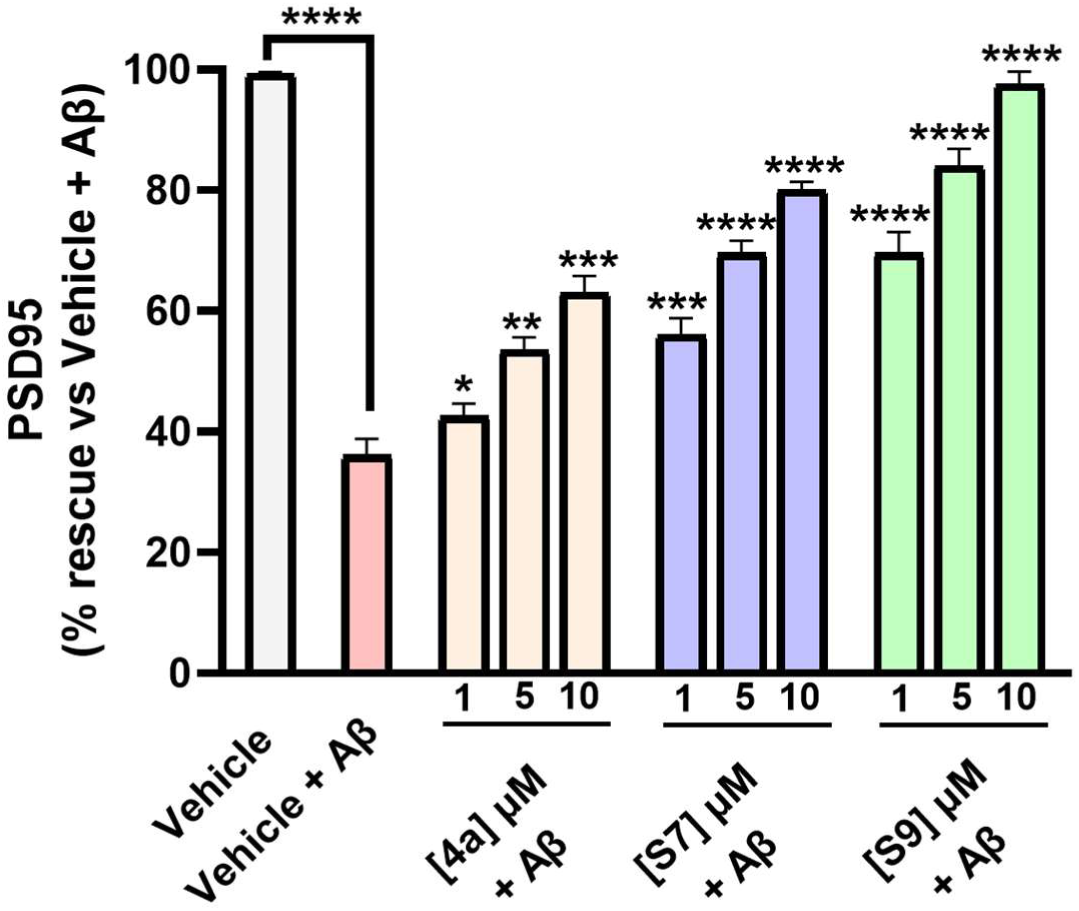
Optimized TREM2 agonists preserve synaptic markers in neuron–microglia co-cultures under amyloid stress. Quantification of PSD95 levels in human iPSC-derived neuron-microglia co-cultures following Aβ₁₋₄₂ oligomer exposure and treatment with **4a**, **S7**, or **S9** (1, 5, and 10 µM). PSD95 levels are expressed as percent rescue relative to Aβ-treated vehicle controls. Data are presented as mean ± SD (n = 5). Statistical significance was assessed by one-way ANOVA with Dunnett’s post hoc test. *p* < 0.05 (*), *p* < 0.01 (**), *p* < 0.001 (***), and *p* < 0.0001 (****).

Taken together, these data demonstrate that second-generation TREM2 agonists **S7** and **S9** exhibit superior functional performance compared with the first-generation lead **4a** across multiple AD-relevant in vitro assays. Importantly, the inclusion of TREM2 KO validation confirms an on-target mechanism of action, while ApoE modulation links compound activity to a core microglial pathway implicated in AD. Among the tested analogs, **S9** consistently emerged as the most promising candidate, combining strong suppression of Aβ-induced inflammatory signaling, TREM2-dependent activity, modulation of ApoE secretion, and preservation of synaptic markers. While the present study focuses on in vitro systems, the convergence of affinity, PK properties, and functional efficacy supports further in vivo evaluation of **S9** as a next-generation small molecule TREM2 agonist.

## 3. Methods

### 3.1. Chemistry

All reactions were performed under protection of N_2_ in oven-dried glassware unless water was applied as solvent. To obtain degassed solvent, a continuous flow of N_2_ was bubbled through the solvent for 3 hours, and a N_2_-filled balloon was attached to maintain an inert atmosphere during solvent withdrawal. The degassed solvent can be stored under nitrogen for extended periods without the need for further purging. Chromatographic purification was performed as flash chromatography with Combi- Flash^®^ Rf+ UV-VIS MS COMP using RediSep^®^ Silver normal phase silica gel columns and solvents indicated as eluent with default pressure. Fractions were collected based on UV absorption at 254 nm and/or 280 nm unless other wavelengths were indicated.

Analytical TLC was performed on Whatman^®^ TLC silica gel UV254 (250 µm) TLC aluminum plates. Visualization was usually accomplished with UV light (254 nm) unless TLC stain was indicated. Proton, carbon and fluorine nuclear magnetic resonance spectra (^1^H NMR, ^13^C NMR and ^19^F NMR) were recorded on a Bruker 500 MHz spectrometer with solvent resonances as the internal standard (^1^H NMR: CDCl_3_ at 7.26 ppm; ^13^C NMR: CDCl_3_ at 77.0 ppm). 1H NMR data are reported as follows: chemical shift (ppm), multiplicity (s = singlet, d = doublet, dd = doublet of doublets, dt = doublet of triplets, ddd = doublet of doublet of doublets, t = triplet, m = multiplet, br = broad), coupling constants (Hz), and integration. Mass spectra were obtained through ESI on Micromass Waters LCT Premier XE. The accurate mass analyses run in EI mode were at a mass resolution of 10,000 to 15,000 FWHM and were calibrated using Leucine Enkephalin acetate as an internal standard.

#### 3.1.2. General method for the synthesis of final product **4m** to **4u**

A 20 mL scintillation vial was charged a stir bar, **3** (106 mg, 0.3 mmol, 1 eq.), boronic acid/ester (0.6 mmol, 2 eq.), Cs_2_CO_3_ (244 mg, 0.75 mmol, 2.5 eq.), and Pd(PPh_3_)_4_ (52 mg, 45 mmol, 15 mol%). A mixture of degassed 1,4-dioxane (2.5 mL) and water (0.5 mL) was added to the vial. The system was purged with N₂ via three cycles of evacuation and backfilling. The reaction mixture was stirred at 85 °C for overnight, cooled to room temperature, and partitioned between DCM (100 mL) and water (100 mL). The organic layer was separated, and the aqueous layer was further extracted with DCM (30 mL). The combined organic layers were washed with brine, dried over anhydrous CaCl_2_ or MgSO_4_, concentrated under reduced pressure to obtain crude product, which was absorbed onto a plug of silica gel and purified by chromatography.

Compound **S1** (6-methoxy-*N*-methyl-5-(3-methylenecyclobutane-1-carboxamido)-[1,1’-biphenyl]- 3-carboxamide). **S1** was obtained (91 mg, 0.260 mmol, 87%) as white color solid by following general procedure with **3** and phenylboronic acid (73 mg), and purification was accomplished by chromatography with 30%-70% EtOAc in hexanes. **^1^H NMR** (500 MHz, CDCl_3_) δ 8.59 (s, 1H), 7.78 (s, 1H), 7.41 – 7.28 (m, 5H), 6.80 (s, 1H), 5.54 (q, *J* = 5.2 Hz, 1H), 4.85 – 4.80 (m, 2H), 3.89 (s, 3H), 3.16 (p, *J* = 8.2 Hz, 1H), 3.12 – 3.05 (m, 2H), 2.95 – 2.90 (m, 2H), 2.69 (d, *J* = 5.0 Hz, 3H). **^13^C NMR** (126 MHz, CDCl_3_) δ 172.59, 169.68, 148.68, 143.96, 140.19, 135.74, 128.54, 128.40, 128.32, 127.45, 126.52, 119.85, 111.58, 106.63, 55.86, 35.90, 35.50, 26.55. **HRMS** m/z: [M+H]^+^ cal. for C21 H23 N2 O3, 351.1709; Found, 351.1711.

Compound **S2** (4-methoxy-*N*-methyl-3-(1-methyl-1*H*-pyrrol-2-yl)-5-(3-methylenecyclobutane-1- carboxamido)benzamide). **S2** was obtained (94 mg, 0.266 mmol, 87%) as yellow color solid by following general procedure with **3** and 1-methyl-2-(4,4,5,5-tetramethyl-1,3,2-dioxaborolan-2-yl)-1*H*-pyrrole (124 mg), and purification was accomplished by chromatography with 30%-70% EtOAc in hexanes. **^1^H NMR** (500 MHz, CDCl_3_) δ 8.87 (s, 1H), 7.70 (s, 1H), 6.80 (s, 1H), 6.72 (s, 1H), 6.20 (s, 1H), 6.15 (s, 1H), 5.46 (s, 1H), 4.84 (d, *J* = 2.4 Hz, 2H), 3.90 (s, 3H), 3.36 (s, 3H), 3.23 – 3.07 (m, 3H), 2.99 – 2.90 (m, 2H), 2.72 (d, *J* = 4.9 Hz, 3H). **^13^C NMR** (126 MHz, CDCl_3_) δ 172.35, 168.19, 148.83, 144.00, 131.70, 129.09, 127.57, 126.40, 123.23, 120.77, 112.92, 108.69, 107.91, 106.72, 55.98, 36.07, 35.60, 34.16, 27.02. **HRMS** m/z: [M+H]^+^ cal. for C20 H24 N3 O3, 354.1818; Found, 354.1816.

Compound **S3** (3-(furan-2-yl)-4-methoxy-*N*-methyl-5-(3-methylenecyclobutane-1- carboxamido)benzamide). **S3** was obtained (91 mg, 0.267 mmol, 89%) as yellow color solid by following general procedure with **3** and furan-2-ylboronic acid (67 mg), and purification was accomplished by chromatography with 30%-70% EtOAc in hexanes. **^1^H NMR** (500 MHz, CDCl_3_) δ 8.40 (s, 1H), 7.80 (s, 1H), 7.41 (d, *J* = 2.0 Hz, 1H), 7.14 (s, 1H), 6.60 (d, *J* = 3.5 Hz, 1H), 6.41 (dd, *J* = 3.4, 1.8 Hz, 1H), 5.97 (q, *J* = 5.1 Hz, 1H), 4.81 (q, *J* = 2.4 Hz, 2H), 3.89 (s, 3H), 3.17 – 3.10 (m, 1H), 3.08 – 3.01 (m, 2H), 2.94 – 2.84 (m, 5H). **^13^C NMR** (126 MHz, CDCl_3_) δ 172.67, 170.31, 151.11, 148.33, 143.90, 141.92, 127.06, 126.51, 123.80, 119.00, 111.83, 108.21, 108.03, 106.65, 55.82, 35.89, 35.47, 26.73. **HRMS** m/z: [M+H]^+^ cal. for C19 H21 N2 O4, 341.1501; Found, 341.1501.

Compound **S5** (4’-chloro-2’-fluoro-6-methoxy-*N*-methyl-5-(3-methylenecyclobutane-1- carboxamido)-[1,1’-biphenyl]-3-carboxamide). **S5** was obtained (102 mg, 0.253 mmol, 84%) as yellow color solid by following general rocedure with **3** and (4-chloro-2-fluorophenyl)boronic acid (105 mg), and purification was accomplished by chromatography with 30%-70% EtOAc in hexanes. **^1^H NMR** (500 MHz, CDCl_3_) δ 8.69 (s, 1H), 7.79 (s, 1H), 7.26 (t, *J* = 8.2, 1H), 7.16 (dd, *J* = 8.2, 2.2 Hz, 1H), 7.11 (dd, *J* = 9.7, 2.1 Hz, 1H), 6.77 (s, 1H), 5.88 (q, *J* = 5.2 Hz, 1H), 4.86 (p, *J* = 2.4 Hz, 2H), 3.91 (s, 3H), 3.22 – 3.15 (m, 1H), 3.14 – 3.07 (m, 2H), 3.00 – 2.90 (m, 2H), 2.82 (d, *J* = 4.9 Hz, 3H). **^13^C NMR** (126 MHz, CDCl_3_) δ 172.89, 168.81, 159.22 (d, *J* = 249.3 Hz), 148.49, 143.75, 134.14 (d, *J* = 10.4 Hz), 131.58 (d, *J* = 4.1 Hz), 129.27 (d, *J* = 10.9 Hz), 129.07, 127.22, 126.97 (d, *J* = 15.4 Hz), 124.40 (d, *J* = 3.6 Hz), 118.65, 116.19 (d, *J* = 25.9 Hz), 112.62, 106.91, 56.00, 36.08, 35.57, 26.67. **HRMS** m/z: [M+H]^+^ cal. for C21 H21 N2 O3 Cl F, 403.1225; Found, 403.1225.

Compound **S6** (2-fluoro-6’-methoxy-*N*^3^*,N*^3^*,N^3^’*-trimethyl-5’-(3-methylenecyclobutane-1- carboxamido)-[1,1’-biphenyl]-3,3’-dicarboxamide). **S6** was obtained (87 mg, 0.198 mmol, 66%) as white color solid by following general procedure with **3** and (3-(dimethylcarbamoyl)-2-fluorophenyl)boronic acid (95 mg, 1.5 eq.), and purification was accomplished by chromatography with 1%-4% MeOH in EtOAc. **^1^H NMR** (500 MHz, CDCl_3_) δ 8.71 (s, 1H), 7.81 (s, 1H), 7.35 (ddt, *J* = 7.9, 4.4, 1.8 Hz, 2H), 7.23 (t, *J* = 7.6 Hz, 1H), 6.80 (s, 1H), 5.93 (q, *J* = 5.0 Hz, 1H), 4.86 (p, *J* = 2.4 Hz, 2H), 3.91 (s, 3H), 3.23 – 3.15 (m, 1H), 3.15 – 3.06 (m, 5H), 3.01 – 2.93 (m, 5H), 2.81 (d, *J* = 4.9 Hz, 3H). **^13^C NMR** (126 MHz, CDCl_3_) δ 172.95, 168.79, 166.83, 154.88 (d, *J* = 246.6 Hz), 148.57, 143.75, 131.79 (d, *J* = 3.2 Hz), 130.00, 129.01, 128.87 (d, *J* = 16.3 Hz), 128.13 (d, *J* = 4.1 Hz), 127.17, 124.56 (d, *J* = 19.5 Hz), 124.53 (d, *J* = 4.1 Hz), 118.41, 112.80, 106.92, 56.05, 38.50 (d, *J* = 3.2 Hz), 36.11, 35.58, 34.93, 26.60. **HRMS** m/z: [M+H]^+^ cal. for C24 H27 N3 O4 F, 440.1986; Found, 440.1982.

Compound **S7** (4-methoxy-*N*-methyl-3-(3-methylenecyclobutane-1-carboxamido)-5-(naphthalen- 1-yl)benzamide). **S7** was obtained (93 mg, 0.232 mmol, 75%) as yellow color solid by following general procedure with **3** and naphthalen-1-ylboronic acid (107 mg), and purification was accomplished by chromatography with 70% EtOAc in hexanes. **^1^H NMR** (500 MHz, CDCl_3_) δ 8.83 (s, 1H), 7.91 – 7.88 (m, 1H), 7.87 (d, *J* = 8.2 Hz, 1H), 7.78 (s, 1H), 7.62 (dd, *J* = 8.4, 0.9 Hz, 1H), 7.53 – 7.46 (m, 2H), 7.41 (td, *J* = 7.8, 1.4 Hz, 2H), 6.83 (s, 1H), 5.37 – 5.28 (m, 1H), 4.86 (p, *J* = 2.4 Hz, 2H), 3.87 (s, 3H), 3.26 – 3.10 (m, 3H), 3.04 – 2.93 (m, 2H), 2.43 (d, *J* = 5.0 Hz, 3H). **^13^C NMR** (126 MHz, CDCl_3_) δ 172.59, 168.56, 148.66, 144.00, 138.33, 134.19, 133.48, 131.92, 129.55, 128.36, 128.14, 127.00, 126.74, 126.47, 125.99, 125.50, 125.27, 120.01, 112.76, 106.77, 55.99, 36.11, 35.65, 35.63, 26.47. **HRMS** m/z: [M+H]^+^ cal. for C25 H25 N2 O3, 401.1865; Found, 401.1865.

Compound **S8** (3-(isoquinolin-8-yl)-4-methoxy-*N*-methyl-5-(3-methylenecyclobutane-1- carboxamido)benzamide). **S8** was obtained (68 mg, 0.169 mmol, 56%) as yellow color solid by following general procedure with **3** and isoquinolin-8-ylboronic acid (104 mg), and purification was accomplished by chromatography with 2%-5% MeOH in DCM. **^1^H NMR** (500 MHz, CDCl_3_) δ 9.02 (s, 1H), 8.82 (s, 1H), 8.51 (d, *J* = 5.8 Hz, 1H), 7.86 (s, 1H), 7.82 (d, *J* = 8.2 Hz, 1H), 7.73 – 7.66 (m, 2H), 7.50 (d, *J* = 7.0 Hz, 1H), 6.86 (d, *J* = 3.7 Hz, 1H), 5.87 – 5.73 (m, 1H), 4.90 – 4.85 (m, 2H), 3.89 (d, *J* = 4.0 Hz, 3H), 3.26 – 3.19 (m, 1H), 3.18 – 3.11 (m, 2H), 2.99 (dddt, *J* = 15.6, 6.6, 4.6, 2.3 Hz, 2H), 2.60 (d, *J* = 4.9 Hz, 3H). **^13^C NMR** (126 MHz, CDCl_3_) δ 172.92, 168.38, 150.62, 148.42, 143.82, 142.77, 139.45, 135.87, 133.07, 129.74, 129.62, 128.08, 127.33, 127.19, 126.24, 120.68, 118.79, 113.16, 106.91, 56.05, 36.15, 35.62, 26.56. **HRMS** m/z: [M+H]^+^ cal. for C24 H24 N3 O3, 402.1818; Found, 402.1819.

Compound **S9** (3-(benzofuran-3-yl)-4-methoxy-*N*-methyl-5-(3-methylenecyclobutane-1- carboxamido)benzamide). **S9** was obtained (94 mg, 0.241 mmol, 80%) as yellow color solid by following general procedure with **3** and benzofuran-3-ylboronic acid (97 mg), and purification was accomplished by chromatography with 30%-70% EtOAc in hexanes. **^1^H NMR** (500 MHz, CDCl_3_) δ 8.70 (s, 1H), 7.83 (s, 1H), 7.78 (s, 1H), 7.54(t, *J* = 6.8, 2H), 7.33 (ddd, *J* = 8.4, 7.2, 1.3 Hz, 1H), 7.26 (t, *J* = 8.4, 1H), 7.00 (s, 1H), 5.72 – 5.65 (m, 1H), 4.86 (p, *J* = 2.4 Hz, 2H), 3.93 (d, *J* = 1.7 Hz, 3H), 3.24 – 3.07 (m, 3H), 3.00 – 2.91 (m, 2H), 2.70 (d, *J* = 4.9 Hz, 3H). **^13^C NMR** (126 MHz, CDCl_3_) δ 172.68, 169.42, 155.08, 148.68, 143.88, 142.81, 129.49, 127.24, 126.96, 124.88, 124.59, 123.01, 120.01, 119.88, 119.68, 111.77, 106.82, 56.01, 36.08, 35.60, 26.70. **HRMS** m/z: [M+H]^+^ cal. for C23 H23 N2 O4, 391.1658; Found, 391.1660.

Compound **S10** (4-methoxy-*N*-methyl-3-(6-(1-methyl-1*H*-pyrazol-4-yl)-3,6-dihydro-2*H*-pyran-4- yl)-5-(3-methylenecyclobutane-1-carboxamido)benzamide). **S10** was obtained (120 mg, 0.275 mmol, 92%) as yellow color solid by following general procedure with **3** and 1-methyl-4-(4-(4,4,5,5-tetramethyl-1,3,2- dioxaborolan-2-yl)-5,6-dihydro-2*H*-pyran-2-yl)-1*H*-pyrazole (174 mg), and purification was accomplished by chromatography with 3%-10% MeOH in EtOAc. **^1^H NMR** (500 MHz, CDCl_3_) δ 8.56 (s, 1H), 7.72 (s, 1H), 7.55 (s, 1H), 7.52 (s, 1H), 6.72 (s, 1H), 6.01 (q, *J* = 5.0 Hz, 1H), 5.77 (dt, *J* = 3.1, 1.8 Hz, 1H), 5.27 (q, *J* = 2.6 Hz, 1H), 4.84 (p, *J* = 2.4 Hz, 2H), 3.97 (dt, *J* = 11.1, 5.5 Hz, 1H), 3.91 (s, 3H), 3.89 (s, 3H), 3.82 (dt, *J* = 11.1, 5.3 Hz, 1H), 3.16 (qd, *J* = 8.4, 6.0 Hz, 1H), 3.12 – 3.05 (m, 1H), 2.98 – 2.90 (m, 5H), 2.40 (dq, *J* = 5.5, 2.7 Hz, 2H). **^13^C NMR** (126 MHz, CDCl_3_) δ 172.74, 169.47, 148.57, 143.82, 138.51, 137.36, 136.93, 129.88, 127.71, 126.35, 126.22, 121.92, 118.90, 110.42, 106.85, 67.65, 61.17, 55.92, 38.93, 36.05, 35.58, 35.56, 29.58, 26.75. **HRMS** m/z: [M+H]^+^ cal. for C24 H29 N4 O4, 437.2189; Found, 437.2191.

### 3.2. Single dose affinity screening

TREM2 binding was assessed using Temperature-Related Intensity Change (TRIC) measurements on a Dianthus NT.23PicoDuo instrument (NanoTemper Technologies, Munich, Germany). Recombinant human TREM2 protein (BioTechne, Minneapolis, MN, USA) was labeled with a RED-tris-NTA 2nd Generation His-Tag Labeling Kit (NanoTemper Technologies) according to the manufacturer’s instructions. Labeled TREM2 was diluted to a final concentration of 10 nM in PBST buffer (154 mM NaCl, 5.6 mM Na₂HPO₄, 1.05 mM KH₂PO₄, 0.005% Tween-20, pH 7.4). Test compounds at 30 μM were incubated with labeled TREM2 for 10 minutes at room temperature prior to measurement. Instrument parameters were set at 85% LED excitation power with picomolar detector sensitivity disabled and a laser on-time of 5 seconds. The negative control consisted of labeled TREM2 protein with DMSO vehicle to establish baseline fluorescence, while compound **4a**, a known TREM2 binder, served as the positive control. All measurements were performed in triplicate. Initial data processing was conducted using Dianthus Analysis software (NanoTemper Technologies), and compounds exhibiting normalized fluorescence (Fnorm) changes greater than five standard deviations from the negative control mean were classified as hits.

### 3.3. Dose-Dependent TREM2 Binding

Binding affinity measurements were performed using Microscale Thermophoresis (MST) on a Monolith NT.115 instrument (NanoTemper Technologies). TREM2 protein labeling was performed as described in section 3.2. Labeled TREM2 (40 nM) was incubated with serially diluted compounds (ranging from 100 μM to 1.5 nM) for 10 minutes at room temperature in PBST buffer. Measurements were performed using a red filter, 100% LED power, and medium MST power. Data were analyzed using MO.Affinity Analysis software (NanoTemper Technologies) and GraphPad Prism 9 (GraphPad Software, San Diego, CA, USA).

### 3.4. Cell-based assays

#### AlphaLISA-Based Measurement of SYK Phosphorylation

Levels of SYK phosphorylation were determined using Phospho-AlphaLISA assays specific for phosphorylation at Tyr525/526. This proximity-based assay utilizes two distinct antibodies: one directed against the phosphorylated tyrosine residues and another recognizing a separate region of the SYK protein. Binding of both antibodies to phosphorylated SYK brings donor and acceptor beads into close spatial proximity, leading to the emission of a luminescent signal. The magnitude of this signal reflects the abundance of phosphorylated SYK, as described by the assay manufacturer (Revvity). To account for variations in protein expression, total SYK levels were assessed using the corresponding AlphaLISA Total SYK assay and used for normalization.

HEK cells stably expressing human TREM2 and DAP12, as well as parental HEK cells lacking TREM2 and DAP12, were plated at 50,000 cells per well in 96-well plates containing 100 μL of DMEM (Gibco) supplemented with 10% fetal bovine serum (FBS; Gibco). Cells were maintained for 24 h at 37 °C under humidified conditions with 5% CO₂. Test compounds were prepared in DMEM at selected concentrations, with DMSO used as the vehicle control. Following serum starvation, the culture medium was replaced with compound-containing medium. After 1 h of incubation, the medium was aspirated, cells were lysed, and AlphaLISA assays were performed according to the manufacturer’s instructions (Revvity, USA).

HEK-hTREM2/DAP12 cells and wild-type HEK cells (lacking TREM2 and DAP12 expression) were seeded at a density of 50,000 cells per well in 96-well plates in a final volume of 100 μL of DMEM (Gibco) supplemented with 10% fetal bovine serum (FBS; Gibco). Cells were incubated for 24 h at 37 °C in a humidified atmosphere containing 5% CO₂.

#### Phagocytosis Assay

Phagocytosis assays were conducted in BV2 cells using green fluorescent latex beads (Sigma, #L1030). Prior to use, beads were opsonized in fetal bovine serum (FBS) for 1 h at 37 °C and subsequently diluted in DMEM to final concentrations of 0.01% (v/v) for beads and 0.05% (v/v) for FBS.

BV2 cells were subjected to serum deprivation and subsequently exposed to compounds **4a, S7, S8**, or **S9** (25 μM for 30 min), with vehicle-treated cells serving as controls. Fluorescent beads were added to the culture medium and incubated for an additional 30 min to allow phagocytosis. Cultures were rinsed three times with ice-cold PBS and fixed using 4% paraformaldehyde (PFA; Thermo Fisher Scientific), followed by immunocytochemical labeling.

Images were captured using a 20× objective. Cell numbers were determined manually based on DAPI-stained nuclei. Cells were scored as phagocytic when IBA1-positive cells contained one or more fluorescent beads.

#### Assessment of TREM2 Agonist Activity in Jurkat NFAT Reporter Cells

Jurkat reporter cells carrying a firefly luciferase gene driven by an NFAT-responsive promoter were engineered to express human TREM2 together with its adaptor protein DAP12. Cells were plated in 96- well plates at a density of 1 × 10⁵ cells per well. After 24 h, the test compounds **4a, S7, S8** or **S9** were added to the cultures at a final concentration of 25 μM.

Cells were incubated for 4 h at 37 °C, after which luciferase activity was assessed using the Bright-Glo™ Luciferase Assay System (Promega). Following a 5 min equilibration at room temperature, luminescent signals were recorded using a Tecan microplate reader.

### 3.5. Evaluation of PK and physicochemical properties

The These experiments were conducted following our previously reported methods.^1^ The evaluations included LogD_7.4_ determination, microsomal stability, and kinetic solubility. Solubility was assessed using UV–visible spectrophotometry.

### 3.6. Preparation of Amyloid-β Oligomers

Amyloid-β_1-42_ (Aβ_1-42_) peptide was dissolved in hexafluoro-2-propanol (HFIP), aliquoted, and dried under a gentle stream of nitrogen. The peptide film was resuspended in dimethyl sulfoxide (DMSO) and diluted in phenol-red-free cell culture medium to induce oligomer formation, followed by incubation at 4 °C for 24 h. Prepared Aβ oligomers were used immediately for cellular assays.

### 3.7. Human iPSC-Derived Microglia Culture and Treatment

Human induced pluripotent stem cell (iPSC)-derived microglia were cultured according to the supplier’s protocol in microglia maintenance medium. Cells were seeded into 96-well plates and allowed to equilibrate for 24 h prior to treatment. Compounds **4a**, **S7**, and **S9** were prepared as DMSO stock solutions and diluted into culture medium to final concentrations of 1, 5, and 10 µM (final DMSO ≤0.1%, n=5). Microglia were pretreated with compounds for 1 h, followed by stimulation with Aβ_1-42_ oligomers. After 24 h, culture supernatants were collected for cytokine analysis. Vehicle-treated cells with and without Aβ stimulation served as controls. WT and KO microglia were treated identically in all experiments.

### 3.8. Measurement of IL-1β Secretion

Interleukin-1β (IL-1β) levels in microglial culture supernatants were quantified using a commercially available human IL-1β ELISA kit (from Abcam, Catalog# ab214025) according to the manufacturer’s instructions. Absorbance was measured using a microplate reader, and cytokine concentrations were calculated from a standard curve. Data were normalized to the Aβ-stimulated vehicle control and expressed as percent change relative to this condition.

### 3.9. Measurement of Secreted ApoE

ApoE levels in microglial culture supernatants were quantified using a commercially available human ApoE ELISA kit (from Abcam, Catalog # ab108813) according to the manufacturer’s instructions. Supernatants were collected 24 h after Aβ stimulation and compound treatment. ApoE concentrations were calculated from a standard curve and normalized to Aβ-treated vehicle controls.

### 3.10. Neuron-Microglia Co-culture Assay

Human iPSC-derived neurons were cultured on poly-D-lysine/laminin-coated plates until mature neuronal networks were established. iPSC-derived microglia were added to neuronal cultures at a neuron-to- microglia ratio of 5:1 and allowed to integrate for 24 h. Co-cultures were treated with **4a**, **S7**, or **S9** (1, 5, or 10 µM, n=5), followed by exposure to Aβ_1-42_ oligomers for 48-72 h.

### 3.11. Quantification of PSD95 Levels

PSD95 levels were quantified using Human PSD95 ELISA Kit from Abcam (Catalog No. 50-269-8625). Data were expressed as percent rescue relative to Aβ-treated vehicle controls and analyzed using GraphPad Prism.

### 3.12. Statistical Analysis

All experiments were performed with a minimum of three independent replicates. Data are presented as mean ± standard deviation (SD). Statistical comparisons between groups were performed using one-way analysis of variance (ANOVA) followed by Dunnett’s post hoc test. Differences were considered statistically significant at *p* < 0.05.

## 4. Conclusions

Herein, we report the structure-guided optimization of a first-generation micromolar TREM2 binder (**4a**) into a series of potent, drug-like small-molecule agonists through strategic replacement of a lipophilic thiophene moiety with polar benzene-based heterocycles. Among the synthesized derivatives, compound S9 emerged as the lead candidate, demonstrating submicromolar binding affinity (K_D_ = 0.95 µM) and representing a 13-fold improvement over the parent compound **4a**. Beyond binding affinity, functional characterization in cell-based assays confirmed that these optimized molecules function as bona fide TREM2 agonists capable of initiating physiologically relevant signaling cascades. Both **S7** and **S9** robustly triggered proximal Syk kinase phosphorylation, activated downstream NFAT-mediated transcriptional programs, and enhanced microglial phagocytic capacity as evidenced by increased APOE internalization. These mechanistic studies established that the structural modifications preserved and enhanced agonistic activity while simultaneously improving pharmaceutical properties.

Comprehensive pharmacokinetic profiling revealed that **S9** possesses a significantly superior developability profile compared to the original hit compound **4a**. Among the key improvements in the pharmacokinetic properties was 7-fold enhancement in aqueous solubility (from 10.6 µM to 74.1 µM in PBS pH 7.4) as well as reduced lipophilicity bringing LogD closer to the optimal range for CNS penetration (from 6.4 to 4.2). Moreover, S9 exhibited substantially improved metabolic stability with 3.4-fold and 2.4- fold longer plasma half-lives in human and mouse respectively, and a 78% reduction in human intrinsic clearance (from 42 to 9 mL/min/mg) when compared to the parent compound **4a**. Critically, **S9** exhibited a 9-fold improved hERG safety margin (IC₅₀ = 28.1 µM vs 3.2 µM for 4a) which substantially mitigates cardiotoxicity risk. Together, the improved pharmacokinetic properties position **S9** as a viable candidate for oral administration and CNS drug development.

To validate therapeutic relevance, we evaluated **S9** in disease-relevant human cellular models. The treatment of **S9** in human iPSC-derived microglia challenged with amyloid-beta oligomers resulted in significant suppression of the secretion of pro-inflammatory cytokine IL-1β through a mechanism that was abolished in TREM2-deficient cells. Furthermore, in human neuron-microglia co-culture systems exposed to amyloid stress, **S9** preserved synaptic integrity as measured by maintenance of postsynaptic density protein 95 (PSD95) expression levels which demonstrates neuroprotective efficacy beyond simply modulating microglial inflammation. Our findings establish that **S9** engages TREM2 in a physiologically meaningful manner that translates to disease-modifying effects in human cellular models of Alzheimer’s pathology. To the best of our knowledge, this work establishes **S9** as a first submicromolar TREM2 agonist with promising oral bioavailability, validated on-target activity, favorable pharmaceutical properties and demonstrated efficacy in human Alzheimer’s disease models.

## Notes

The authors declare no competing financial interests.

