## Supporting Information for "Small Molecule Agonists of TREM2 Reprogram Microglia and Protect Synapses in Human Alzheimer’s Models"

Corresponding:

^*^Moustafa T. Gabr: Department of Radiology, Molecular Imaging Innovations Institute (MI3), Weill Cornell Medicine, New York, NY 10065, USA.

**Table of Content**

I. Supplementary figures	[S3](#_Toc189481170)

II. [Spectral data	S4](#_Toc189481172)

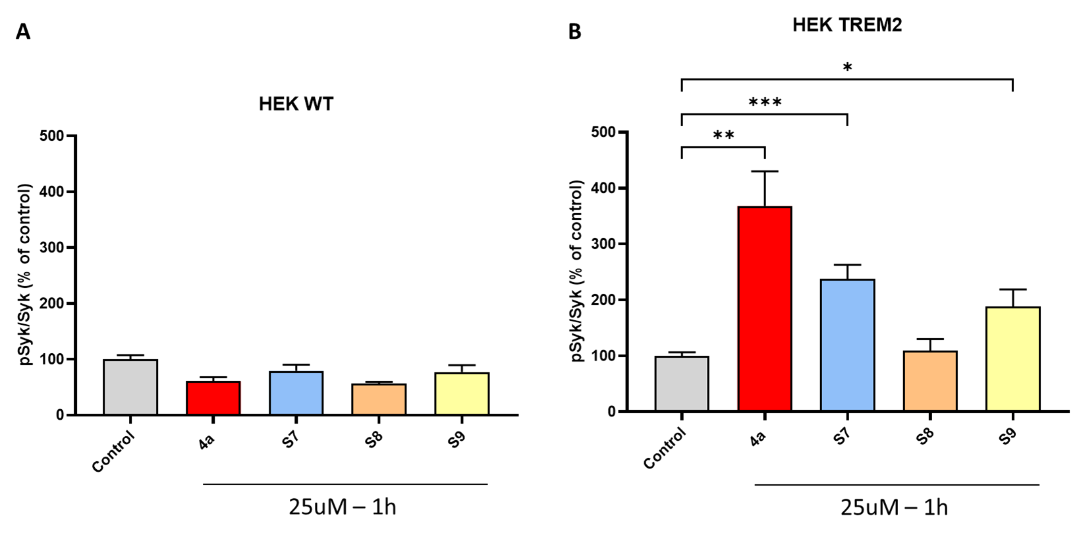

**Figure S1. In vitro assessment of TREM2-mediated SYK phosphorylation.** (A) Histogram showing phospho-SYK levels quantified by AlphaLISA in HEK cells either expressing TREM2/DAP12 or lacking these proteins, under untreated conditions or following treatment with compounds **4a**, **S7**, **S8**, or **S9** (25 μM). Phospho-SYK levels in treated samples are expressed as a percentage of the untreated control (n = 5 biological replicates). No significant differences in SYK phosphorylation were detected among any of the conditions in the cells lacking TREM2/DAP12.

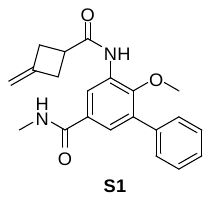

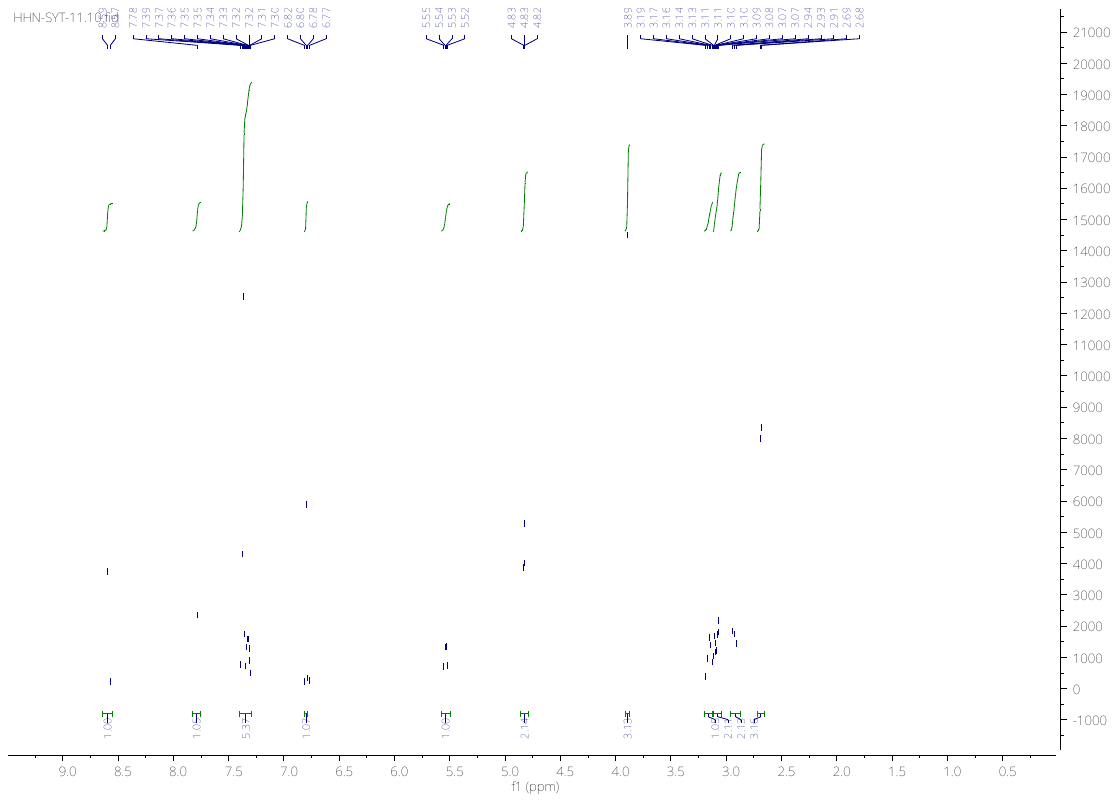
**Spectral data for the synthesized derivatives**

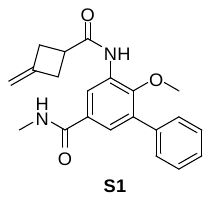

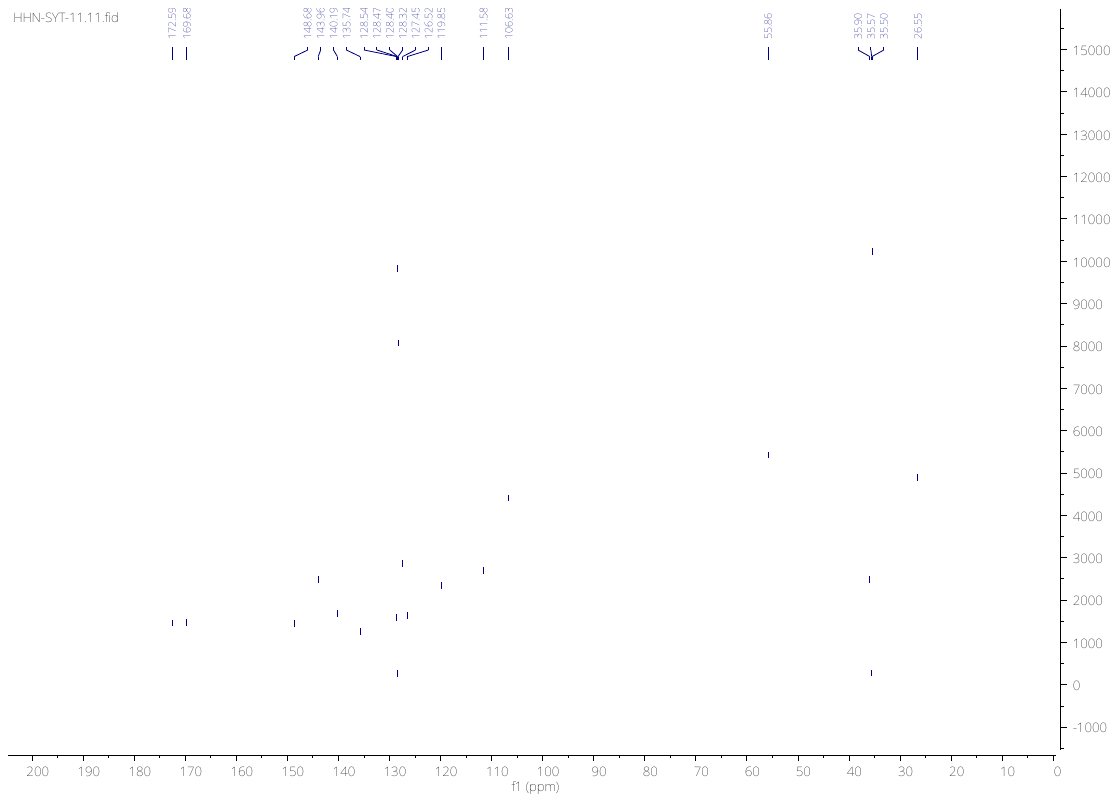

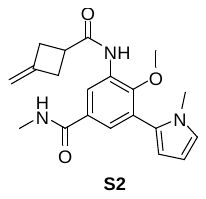

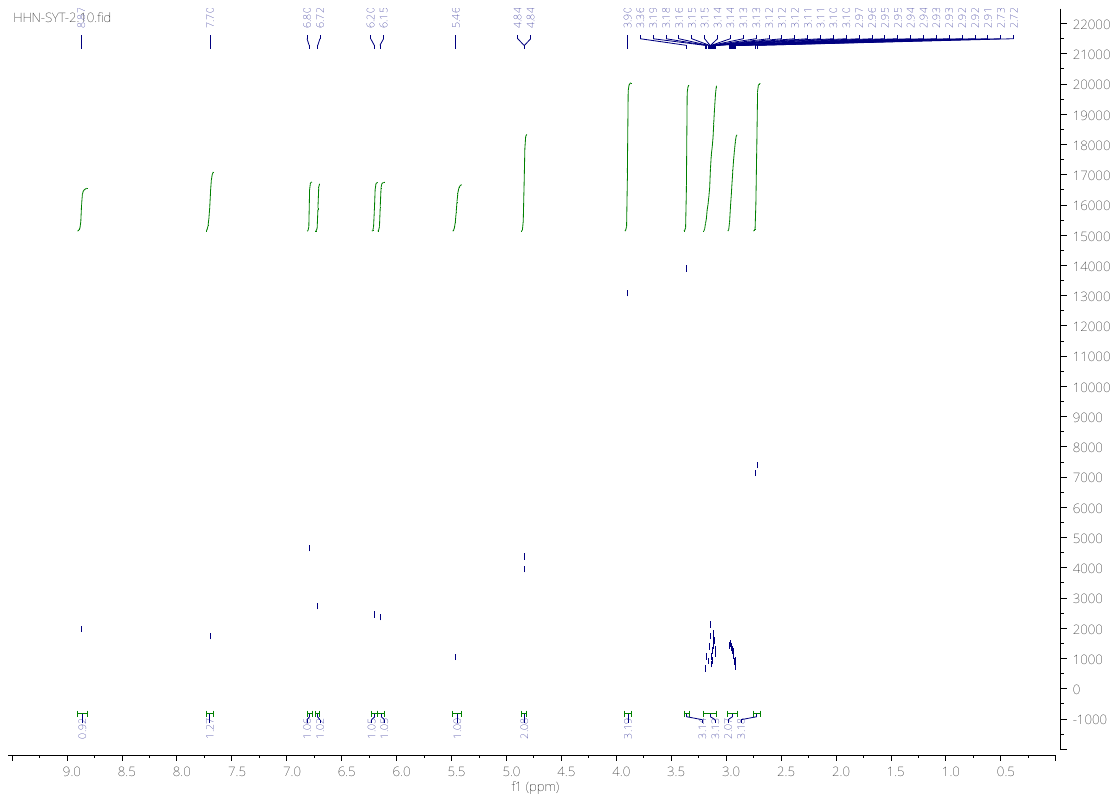

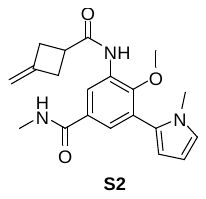

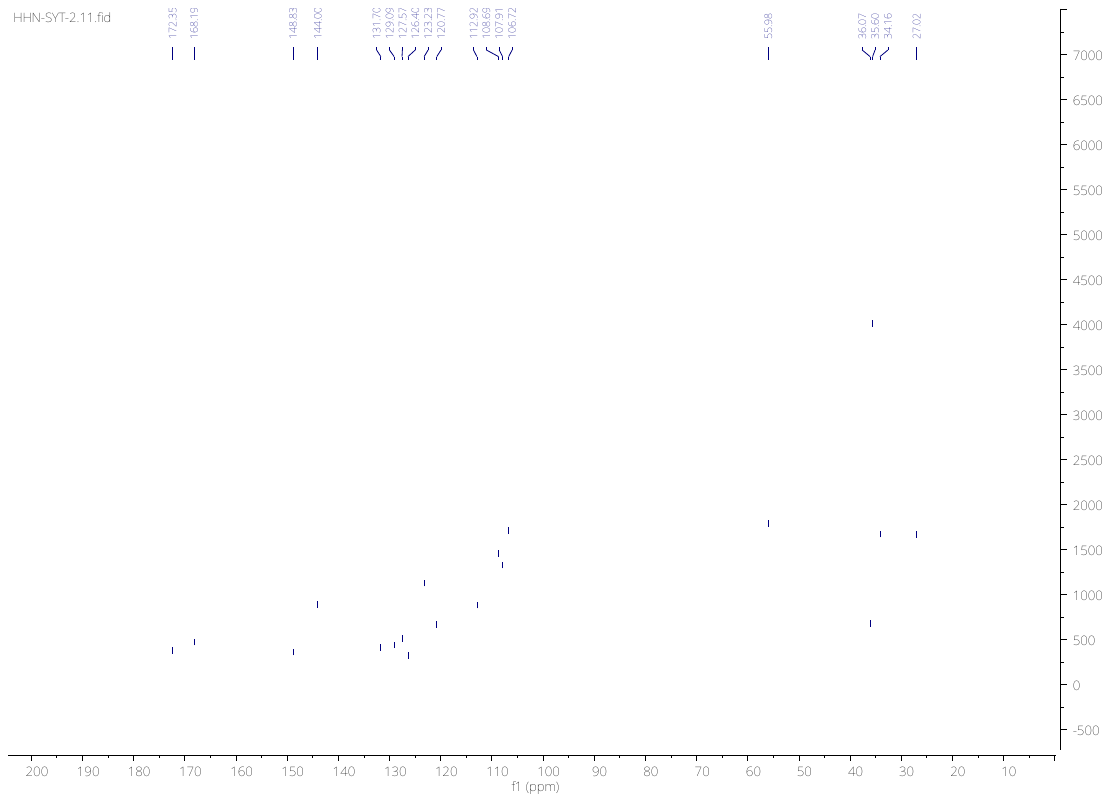

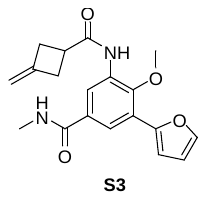

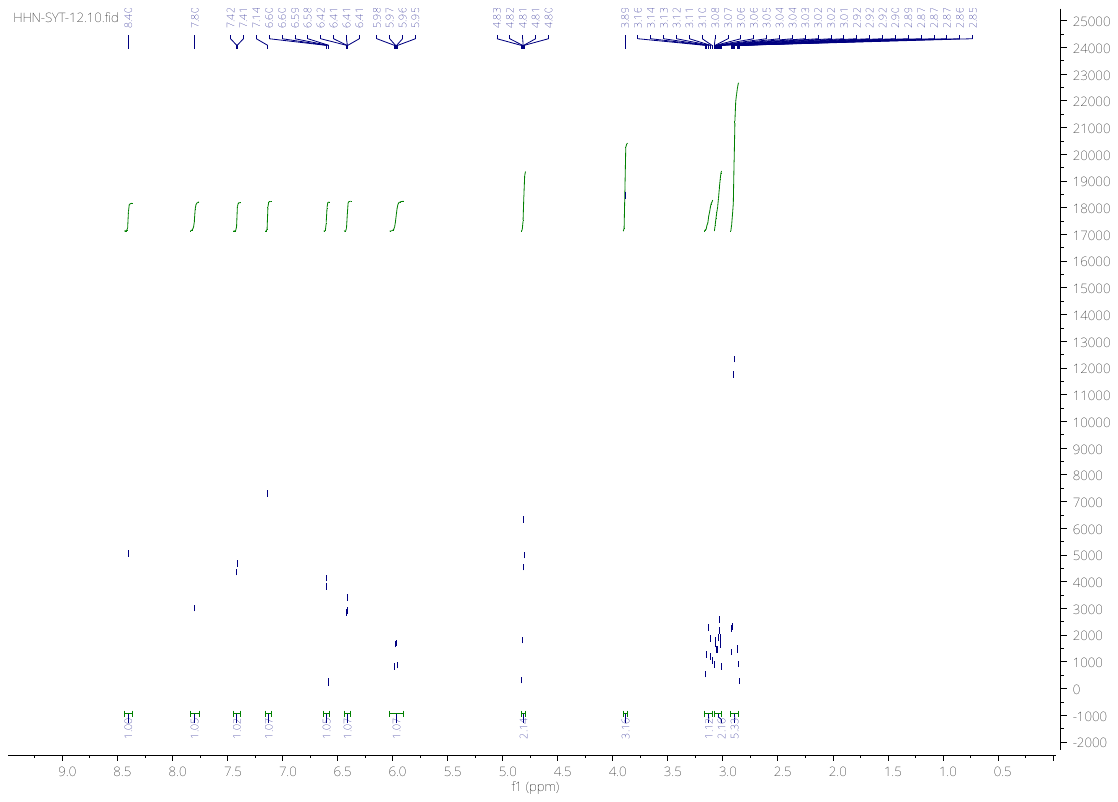

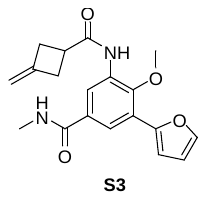

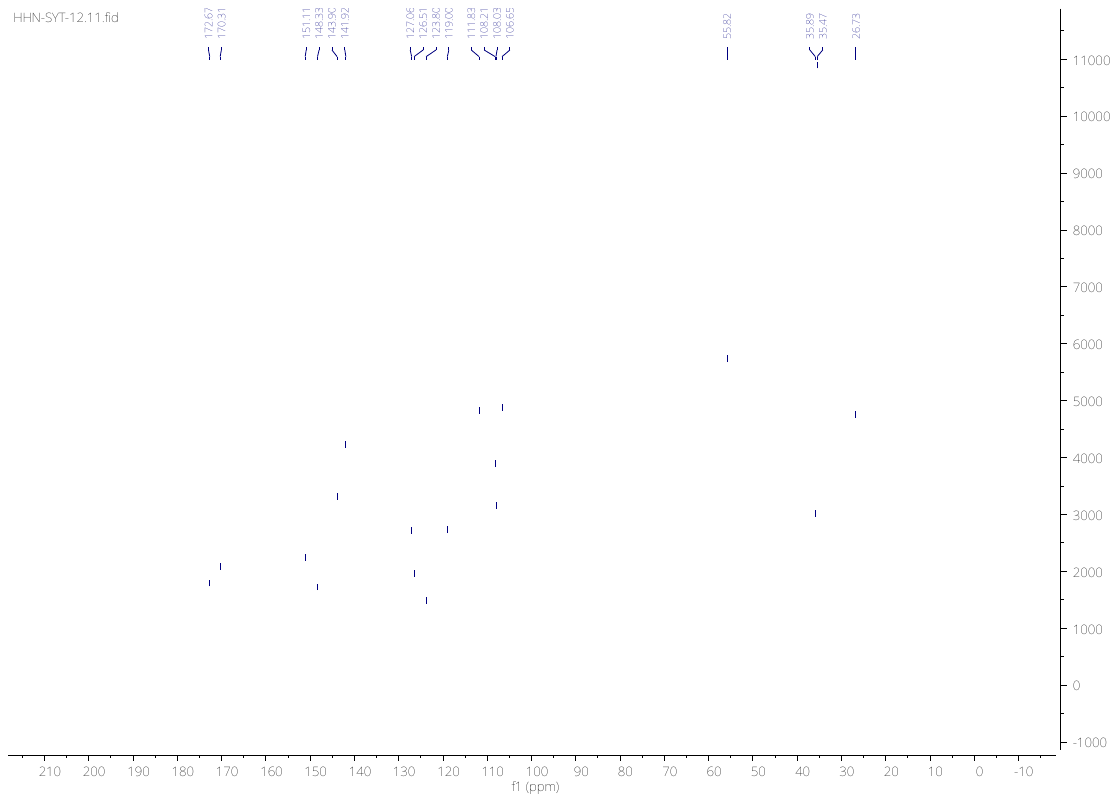

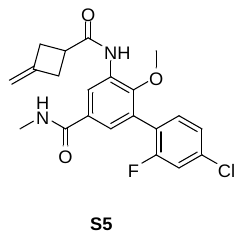

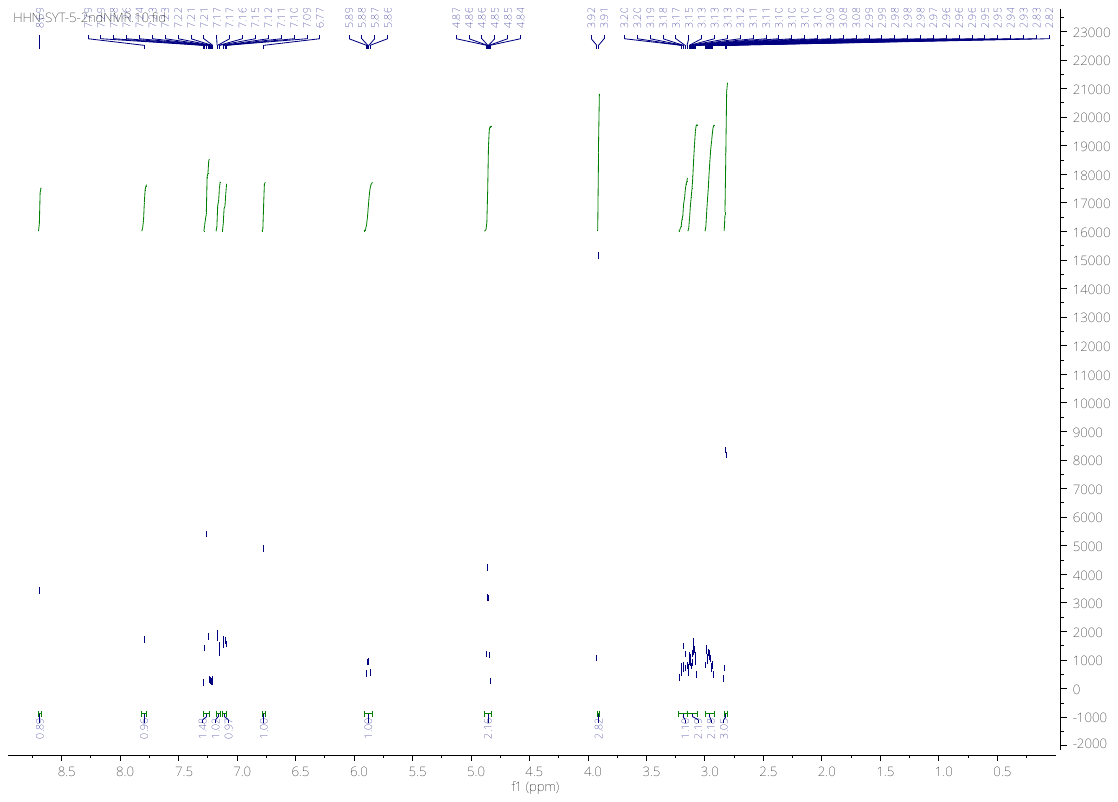

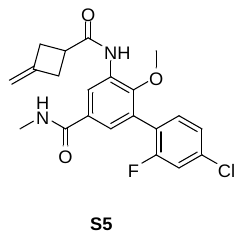

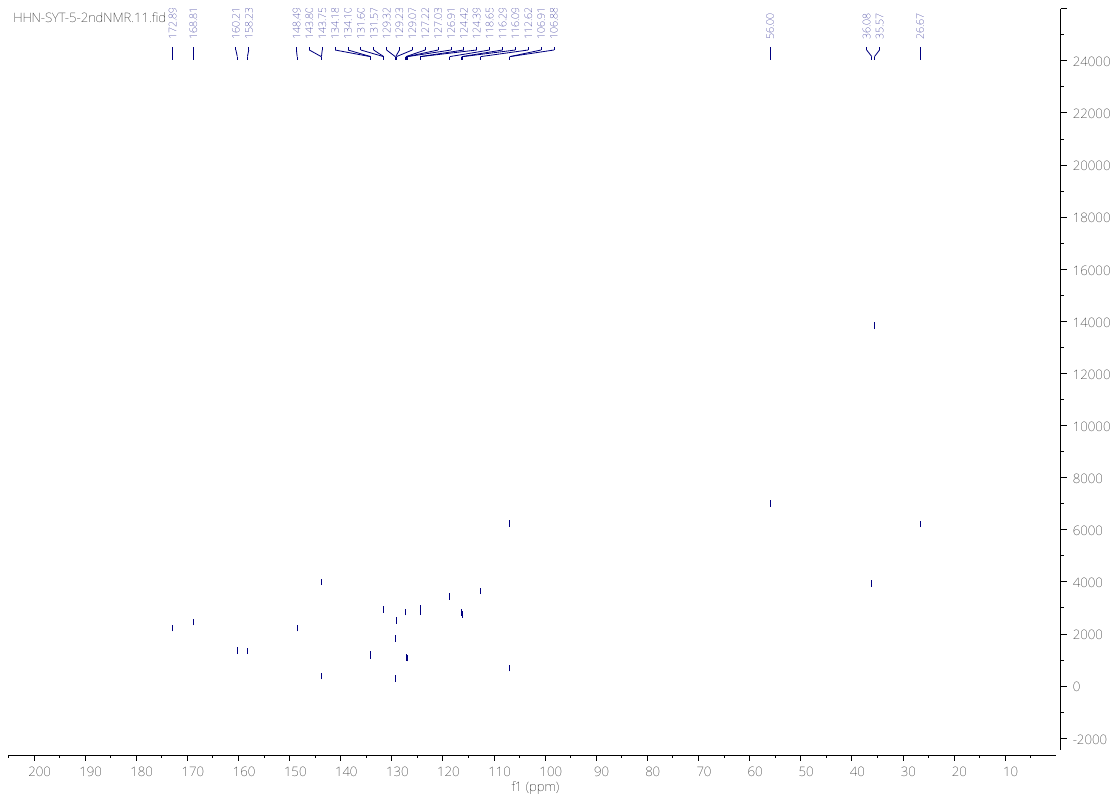

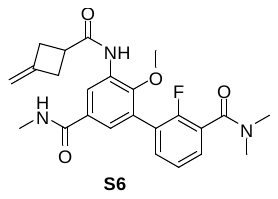

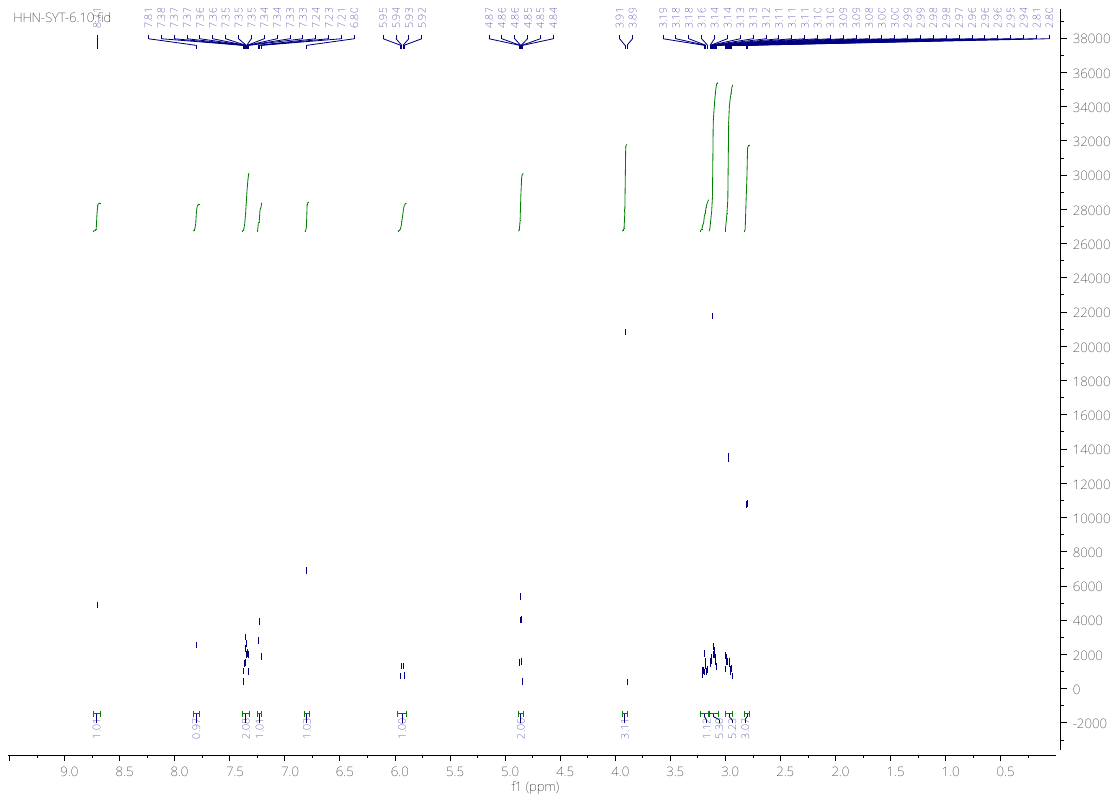

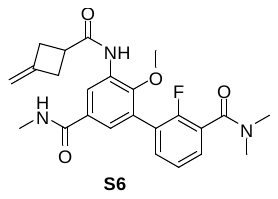

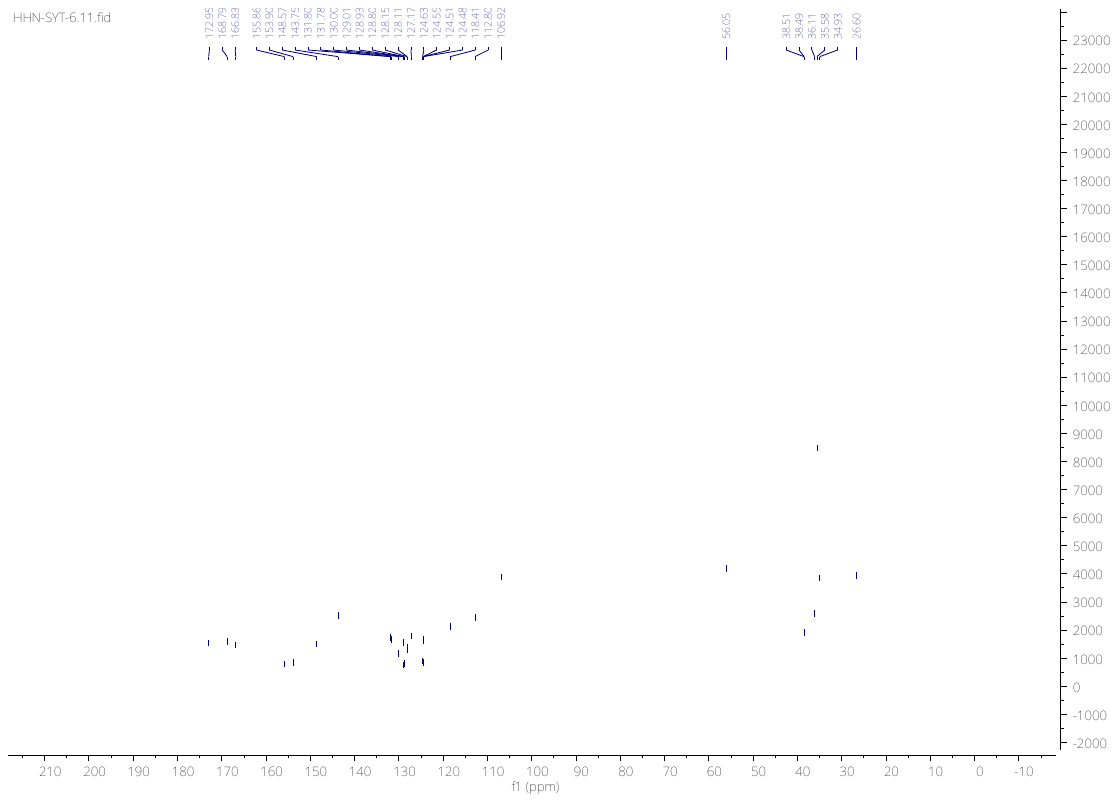

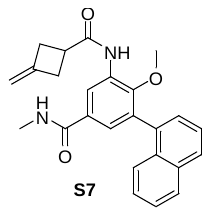

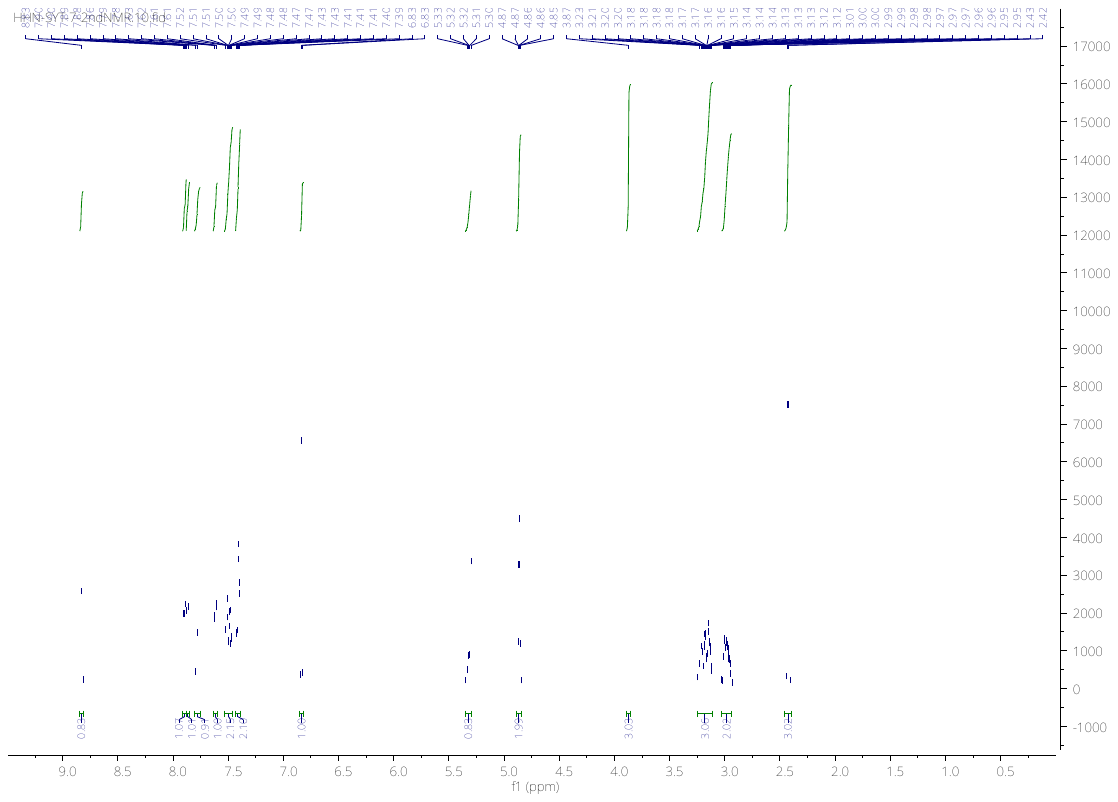

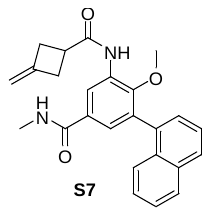

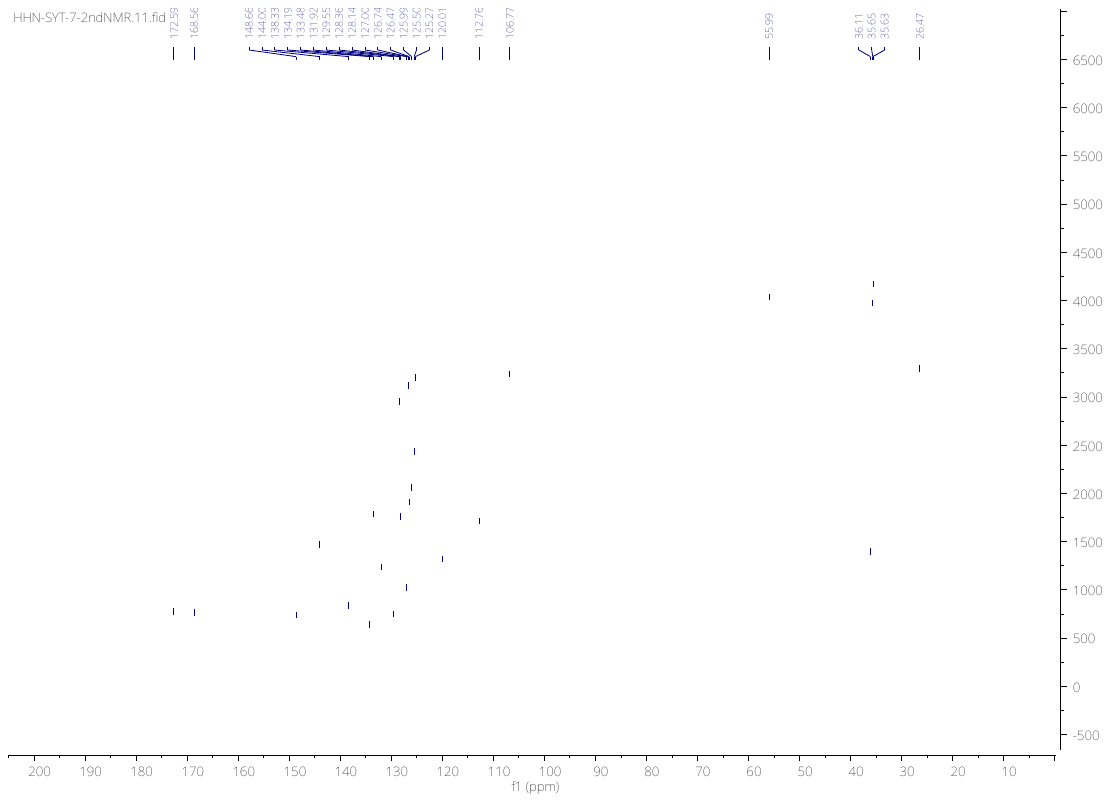

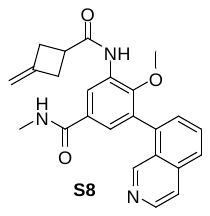

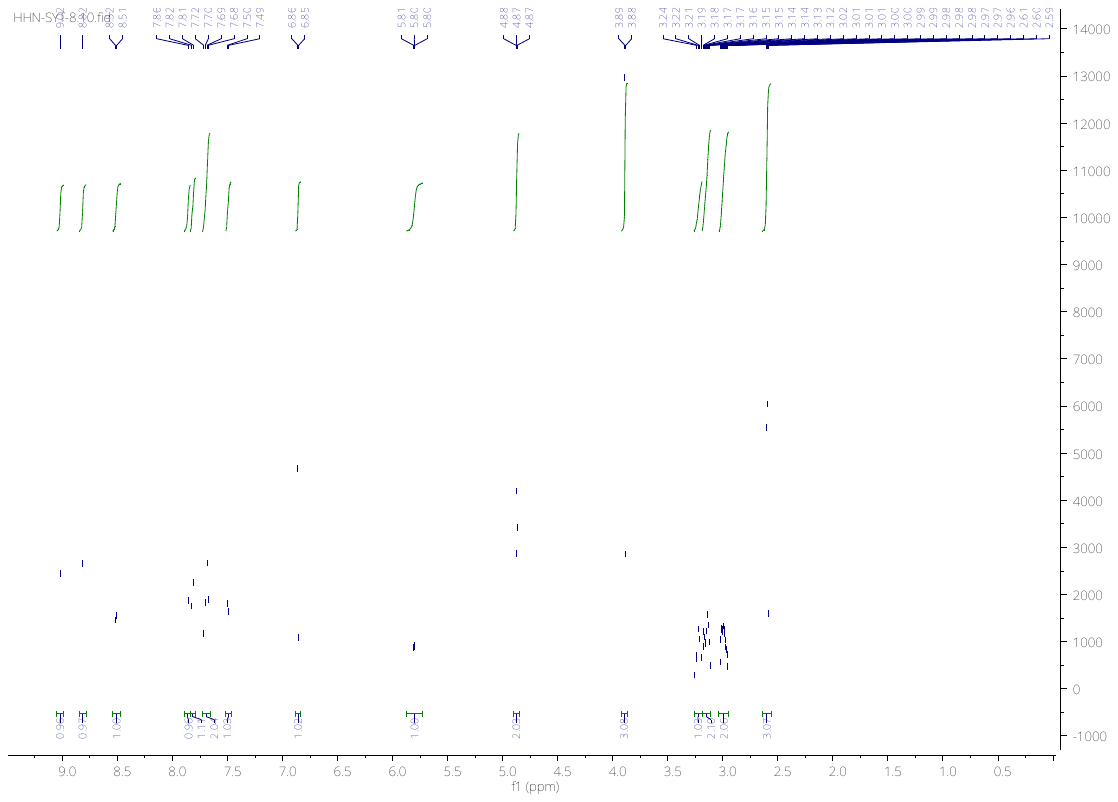

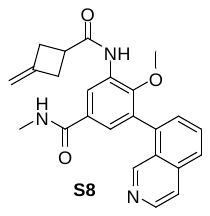

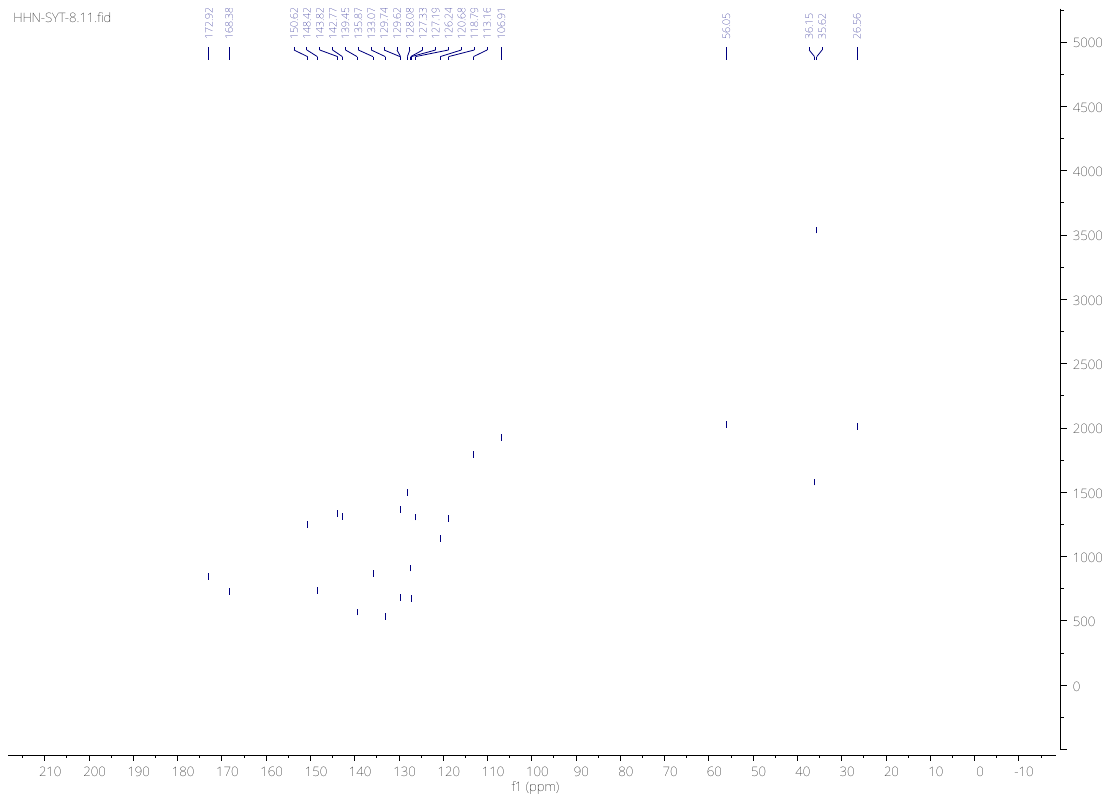

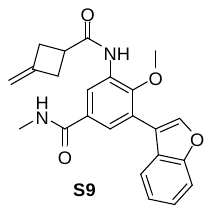

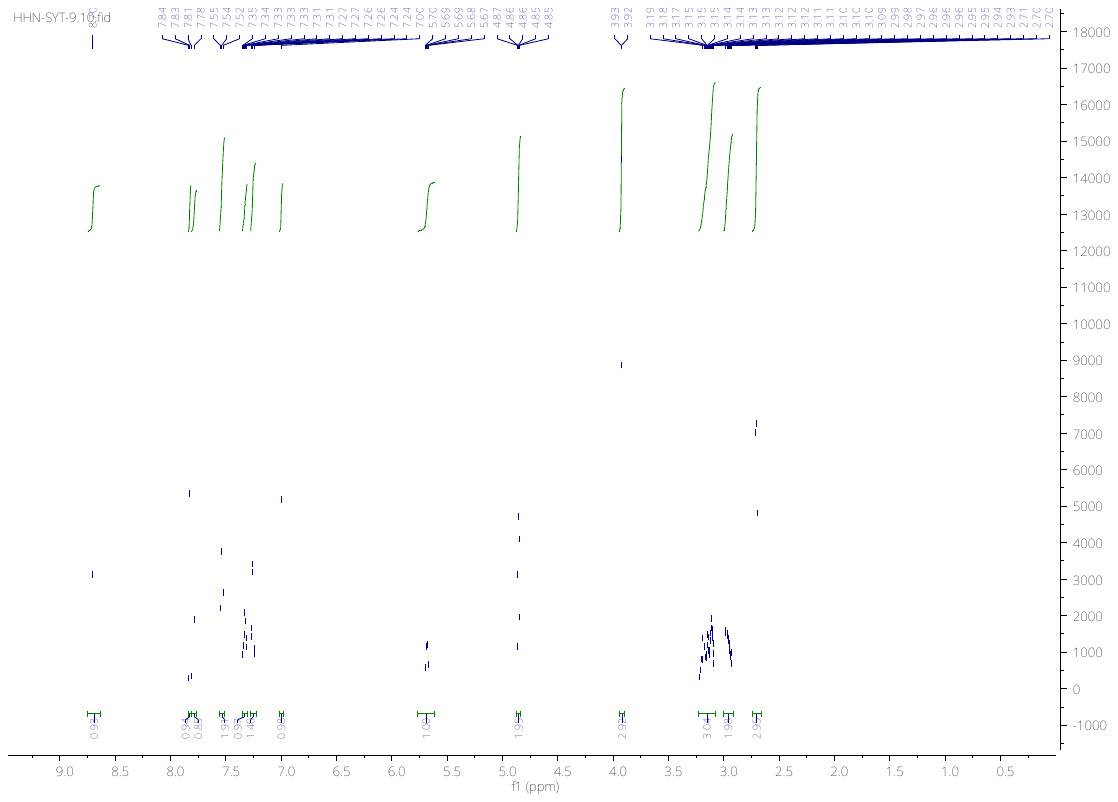

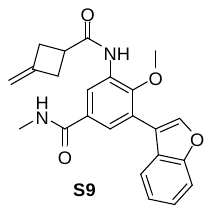

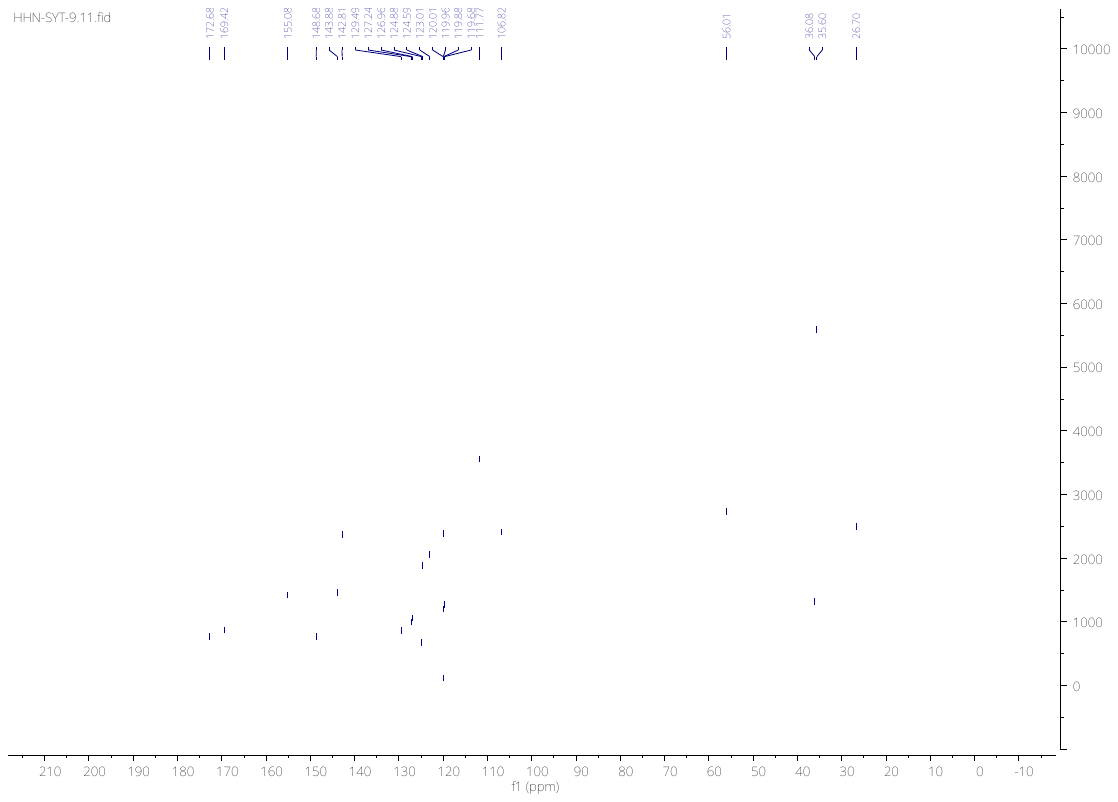

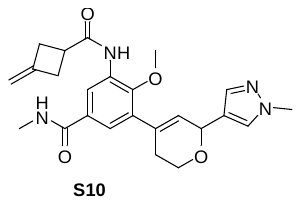

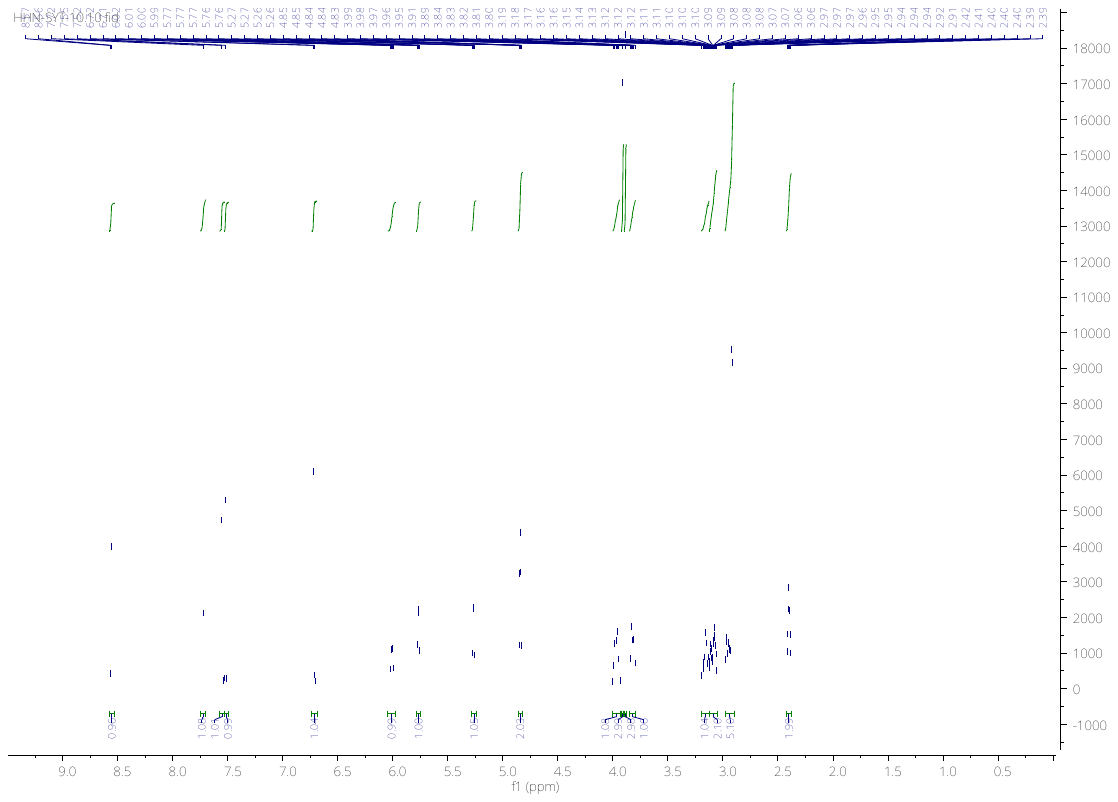

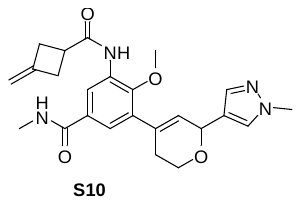

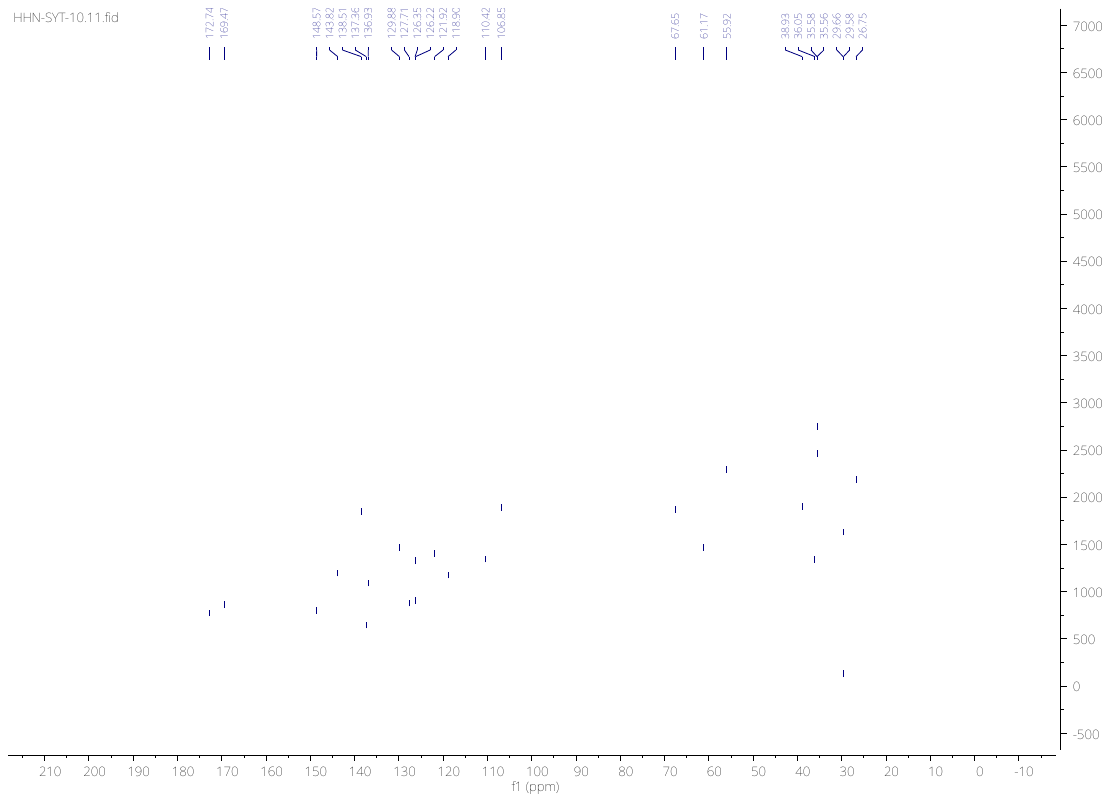

100

351.1711

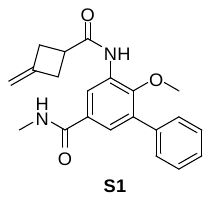

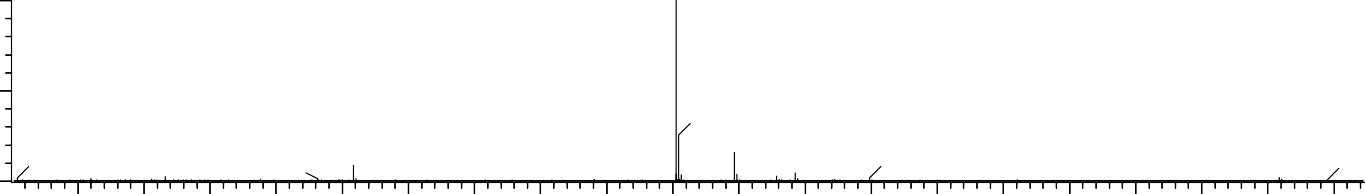

352.1742

102.0135 158.0032 215.5759 229.1416

279.0939 320.1288

396.2291

424.2603 480.3232

545.2253 579.3052 597.3304

m/z

%

0

125	150	175	200	225	250	275	300	325	350	375	400	425	450	475	500	525	550	575	600

| Minimum: |  |  |  | -1.5 |  | | |
| --- | --- | --- | --- | --- | --- | --- | --- |
| Maximum: |  | 5.0 | 5.0 | 50.0 |  |  |  |
| Mass | Calc. Mass | mDa | PPM | DBE | i-FIT | Norm | Conf(%) Formula |
| 351.1711 | 351.1709 | 0.2 | 0.6 | 11.5 | 1284.6 | n/a | n/a C21 H23 N2 O3 |

100

354.1816

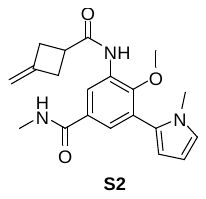

%

0

100	125	150	175	200	225	250	275	300	325	350	375	400	425	450	475	500	525	550	575	600

m/z

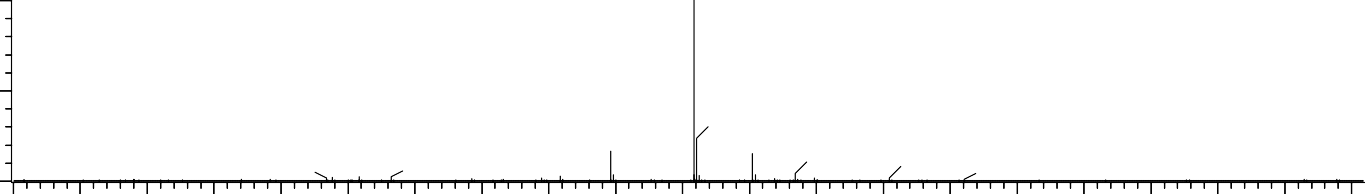

355.1848

144.9825	217.0812229.1414241.1779

323.1394

304.3004

392.1377 427.2709

455.3019

538.2896	582.3157

| Minimum: |  |  |  | -1.5 |  | | |
| --- | --- | --- | --- | --- | --- | --- | --- |
| Maximum: |  | 5.0 | 5.0 | 50.0 |  |  |  |
| Mass | Calc. Mass | mDa | PPM | DBE | i-FIT | Norm | Conf(%) Formula |
| 354.1816 | 354.1818 | -0.2 | -0.6 | 10.5 | 1694.5 | n/a | n/a C20 H24 N3 O3 |

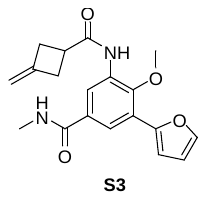

# C: 0-25 H: 0-29 N: 0-4 O: 0-4

C19H20N2O4

HHN_SYT_12 41 (0.182) Cm (41:52)	1: TOF MS ES+

1.36e+008

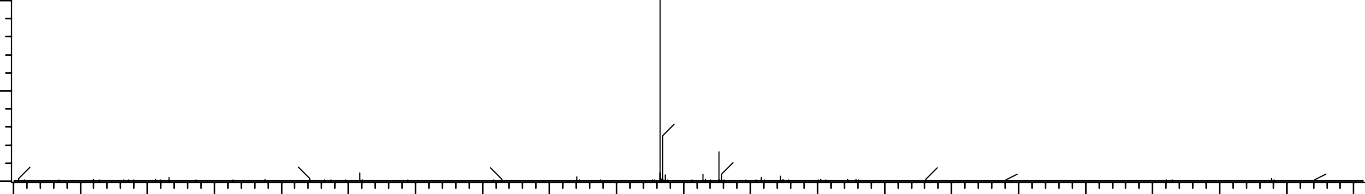

342.1533

102.0134 158.0032 210.5653 229.1415 282.2798

310.1079

364.1352

414.2393

440.2550 470.3022

532.2040 569.2840 585.2481

m/z

100

341.1501

%

0

100	125	150	175	200	225	250	275	300	325	350	375	400	425	450	475	500	525	550	575	600

| Minimum: |  |  |  | -1.5 |  | | |
| --- | --- | --- | --- | --- | --- | --- | --- |
| Maximum: |  | 5.0 | 5.0 | 50.0 |  |  |  |
| Mass | Calc. Mass | mDa | PPM | DBE | i-FIT | Norm | Conf(%) Formula |
| 341.1501 | 341.1501 | 0.0 | 0.0 | 10.5 | 1465.9 | n/a | n/a C19 H21 N2 O4 |

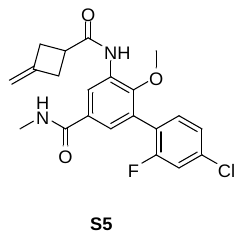
C: 0-21	H: 0-21	N: 0-3	O: 0-3	Cl: 0-1	F: 0-1

C21H20ClFN2O3

HHN_SYT_5 48 (0.209) Cm (48:61)	1: TOF MS ES+

1.64e+008

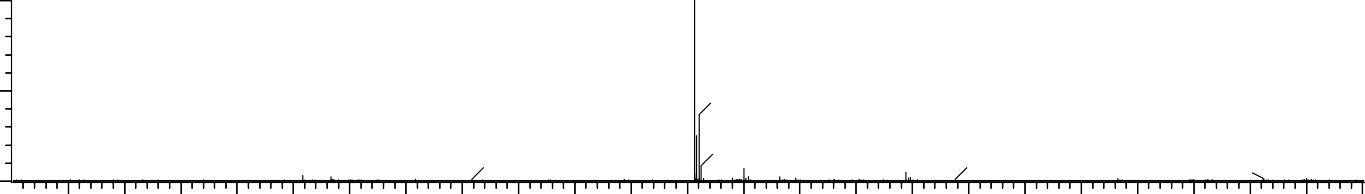

405.1202

158.0030

229.1414

279.0935 304.3005

406.1231

372.0802

497.1443 519.1268

591.1664 656.0899 675.0847

100

403.1225

%

0	m/z

|  | 150 | 200 | 250 |  | 300 | 350 | 400 |  | 450 | 500 | 550 | 600 | 650 | 700 |
| --- | --- | --- | --- | --- | --- | --- | --- | --- | --- | --- | --- | --- | --- | --- |
| Minimum: |  |  |  |  |  | -1.5 |  |  |  |  |  |  |  |  |
| Maximum: |  |  | 5.0 | 5.0 |  | 50.0 |  |  |  |  |  |  |  |  |
| Mass | Calc. | Mass | mDa | PPM |  | DBE | i-FIT | Norm |  | Conf(%) | Formula |  |  |  |
| 403.1225 | 403.1225 | | 0.0 | 0.0 | 11.5 | | 1348.0 | n/a | n/a | | C21 H21 | N2 O3 Cl F | | |

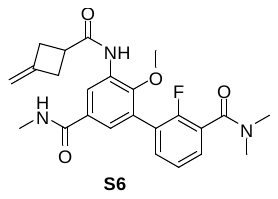

# C: 0-24 H: 0-27 N: 0-3 O: 0-4 F: 0-1

C24H26FN3O4

HHN_SYT_6 59 (0.251) Cm (59:66)	1: TOF MS ES+

3.19e+007

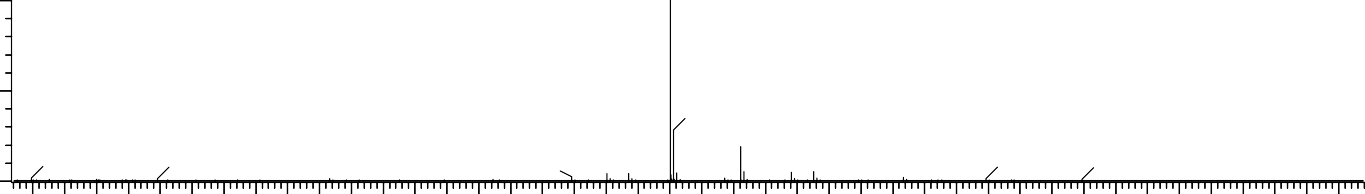

441.2014

239.5762	279.0935

333.1426

409.1563 420.1924

485.2565 513.2878	539.3035 569.3505 614.4851

648.6299

100

440.1982

%

0

240	260	280	300	320	340	360	380	400	420	440	460	480	500	520	540	560	580	600	620	640

m/z

| Minimum: |  |  |  | -1.5 |  | | |
| --- | --- | --- | --- | --- | --- | --- | --- |
| Maximum: |  | 5.0 | 5.0 | 50.0 |  |  |  |
| Mass | Calc. Mass | mDa | PPM | DBE | i-FIT | Norm | Conf(%) Formula |
| 440.1982 | 440.1986 | -0.4 | -0.9 | 12.5 | 1122.8 | n/a | n/a C24 H27 N3 O4 F |

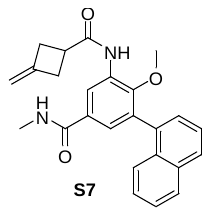
C25H24N2O3

HHN_SYT_7 66 (0.289) Cm (66:72)	1: TOF MS ES+

2.14e+007

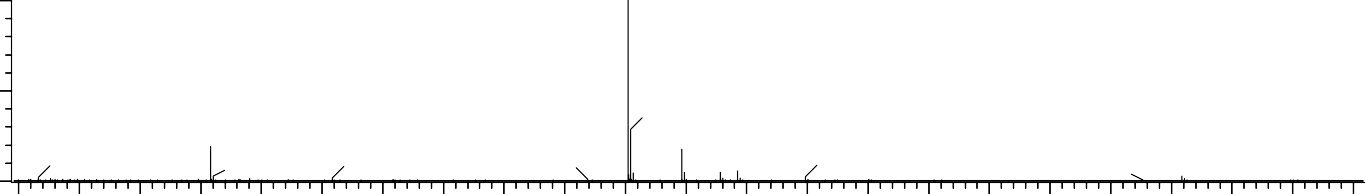

402.1899

158.0030

229.1414

230.1449

279.0938

384.3844

446.2446

474.2759 530.3383

613.3263 629.3209 675.0852

100

401.1865

%

0

150 175 200 225 250 275 300 325 350 375 400 425 450 475 500 525 550 575 600 625 650 675 700

m/z

| Minimum: |  |  |  | -1.5 |  | | |
| --- | --- | --- | --- | --- | --- | --- | --- |
| Maximum: |  | 5.0 | 5.0 | 50.0 |  |  |  |
| Mass | Calc. Mass | mDa | PPM | DBE | i-FIT | Norm | Conf(%) Formula |
| 401.1865 | 401.1865 | 0.0 | 0.0 | 14.5 | 1095.6 | n/a | n/a C25 H25 N2 O3 |

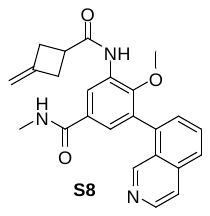
C24H23N3O3

HHN_SYT_8 65 (0.273) Cm (65:67)	1: TOF MS ES+

1.06e+007

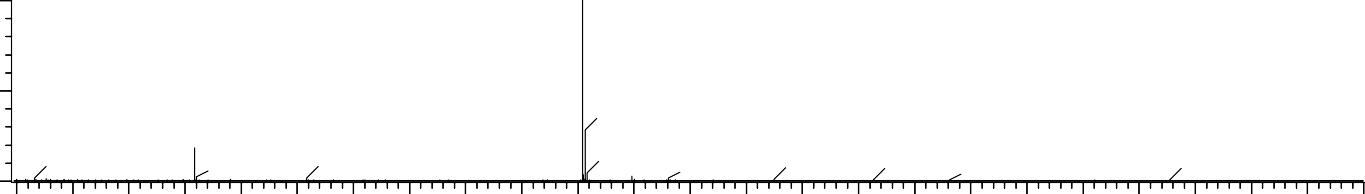

%

403.1852

229.1416

158.0031	230.1450 279.0939 338.3425

404.1880	487.3611 531.3878	663.4575

440.1382	565.5687 630.3161

0

743.5477

m

100

402.1819

/z

| 150 | 200 | 250 | 300 |  | 350 | 400 | 450 |  | 500 | 550 | 600 | 650 | 700 | 750 |
| --- | --- | --- | --- | --- | --- | --- | --- | --- | --- | --- | --- | --- | --- | --- |
| Minimum: |  |  |  |  |  | -1.5 |  |  |  |  |  |  |  |  |
| Maximum: |  |  | 5.0 | 5.0 |  | 50.0 |  |  |  |  |  |  |  |  |
| Mass | Calc. | Mass | mDa | PPM |  | DBE | i-FIT | Norm |  | Conf(%) | Formula |  |  |  |
| 402.1819 | 402.1818 | | 0.1 | 0.2 | 14.5 | | 877.6 | n/a | n/a | | C24 H24 N3 O3 | | | |

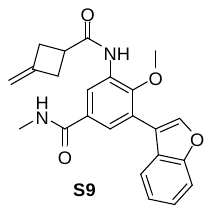
C23H22N2O4

HHN_SYT_9 58 (0.247) Cm (58:65)	1: TOF MS ES+

3.37e+007

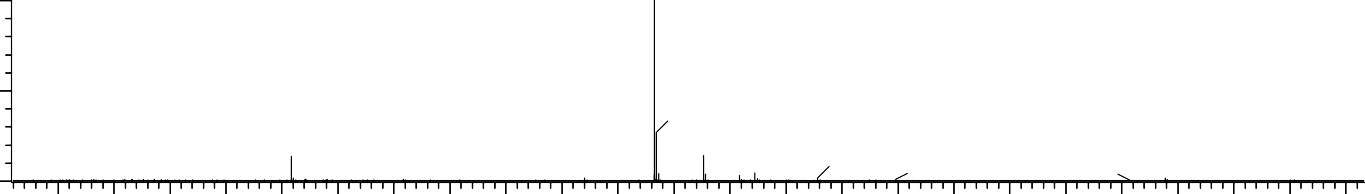

392.1690

229.1415

158.0031

279.0939

360.1238

436.2239

464.2552 499.0991

603.3057 619.2999	675.0851

100

391.1660

%

0

150	200	250	300	350	400	450	500	550	600	650	700

m/z

```
Minimum:	-1.5
Maximum:	5.0	5.0	50.0
Mass	Calc. Mass mDa	PPM	DBE	i-FIT	Norm	Conf(%) Formula 391.1660	391.1658	0.2	0.5	13.5	1136.3 n/a	n/a	C23 H23 N2 O4
```

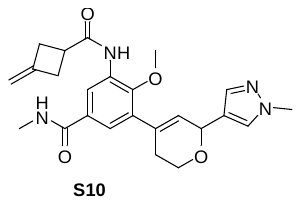
C24H28N4O4

HHN_SYT_10 61 (0.258) Cm (61:65)	1: TOF MS ES+

1.21e+007

100

437.2191

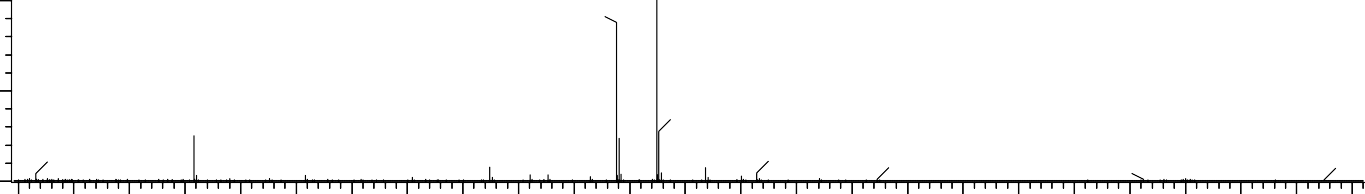
419.2086

%

158.0032

0

229.1415

279.0941

362.1870

438.2224

459.2011 482.2772

536.3243	656.0905 675.0853 715.3051 737.2871

m/z

150	200	250	300	350	400	450	500	550	600	650	700	750

```
Minimum:	-1.5
Maximum:	5.0	5.0	50.0
Mass	Calc. Mass mDa	PPM	DBE	i-FIT	Norm	Conf(%) Formula 437.2191	437.2189	0.2	0.5	12.5	950.7	n/a	n/a	C24 H29 N4 O4
```
